# Slit regulates subcellular-specific targeting of dendrites and axons in the *Drosophila* brain

**DOI:** 10.1101/2024.10.29.620851

**Authors:** Xiaobing Deng, Prachi Shah, Isaac Cervantes-Sandoval, Hui-Hao Lin, Sijun Zhu

## Abstract

Proper nervous system functioning requires axons or dendrites targeting to specific subcellular domains of target neurons to form precise connections. Using mushroom body output neurons (MBONs) and dopaminergic neurons (DAN) in the *Drosophila* mushroom body as a model, we demonstrate that subcellular-specific targeting of MBON dendrites and DAN axons is regulated by repulsion between their dendrites/axons in neighboring subcellular domains. We further show that Slit mediates such repulsion by binding to distinct Robo receptors in different neurons. Loss of Slit-mediated repulsion leads to expansion of MBON dendrites and DAN axons into neighboring compartments, formation of ectopic synapses, and altered learning and memory. Our findings reveal a critical role of Slit-mediated repulsion between neighboring compartments in regulating subcellular-specific dendrite and axon targeting.

## INTRODUCTION

Proper functioning of the nervous system requires neurons to form synapses not only with correct target cells but also at specific subcellular domains (Yogev and Shen 2014). For example, different types of inhibitory GABAergic interneurons in the cerebral cortex form synapses at different subcellular domains with excitatory pyramidal neurons. Parvalbumin-positive basket cells project their axons to the perisomatic region of pyramidal neurons (Klausberger et al. 2003), whereas somatostatin-positive interneurons project their axons to distal dendrites (Kawaguchi and Kubota 1997). Similarly, in the *Drosophila* ventral nerve cord, different sensory neurons project their axons to either medial or lateral dendrites of interneurons to form synapses (Sales et al. 2019; Galindo et al. 2023). The subcellular specificity of synaptic contacts has profound impacts on neuronal activity and computational power of neural circuits (Yogev and Shen 2014).

The subcellular specificity of synaptic contacts can be achieved by targeting of axons to subcellular domains of target neurons or by targeting dendrites to subcellular domain of presynaptic neurons. The mechanisms regulating subcellular-specific targeting of axons can involve both target cells and non-target cells. Target cells express cell surface molecules, such as cell adhesion molecules, receptors, growth factors, at specific subcellular domains to guide axon projection (Ango et al. 2004; Fazzari et al. 2010; Sando et al. 2019). Non-target cells, including neighboring guidepost cells or remote organizers, regulate targeting of axons by expressing non-diffusible or diffusible guidance molecules (Colon-Ramos et al. 2007; Ango et al. 2008; Sales et al. 2019). However, the role of non-target cells in regulating subcellular specificity of synaptic contacts is less well studied, and many of their guidance molecules remain unidentified.

However, subcellular-specific targeting of dendrites has rarely been studied. In the *Drosophila* mushroom body (MB), the olfactory associative learning and memory center (Cognigni et al. 2018), mushroom body output neurons (MBONs) project their dendrites to specific compartments of the MB axonal lobes to form synaptic contacts. MBONs receive synaptic inputs from MB neurons (Kenyon cells, KCs) and mediate behavioral outputs (Cognigni et al. 2018). In each brain hemisphere, there are 34 MBONs of 21 types, each of which project their dendrites to 1-2 of non-overlapping15 compartments in MB axonal lobes (including 5 in the γ lobe, 3 each in the α’ and α lobes, and 2 each in the β’and β lobes) (Fig. 1A) (Tanaka et al. 2008; Aso et al. 2014a). In addition, over 100 local dopaminergic interneurons (DANs) of 20 different types project their axons to 1-2 compartments in the MB axonal lobes (Tanaka et al. 2008; Aso et al. 2014a). Within each compartment, DAN axons form synapses with both MB axons and MBON dendrites, modulating synaptic transmission from MB neurons to MBONs and behavioral outputs in response to unconditioned stimuli (Liu et al. 2012; Aso et al. 2014a; Cohn et al. 2015). Importantly, proper behavioral outputs depend on precise compartmental matching between DAN axons and MBON dendrites. DANs responding to appetitive stimuli typically connect with MBONs that mediate avoidance behavior, while DANs responding to aversive stimuli connect with MBONs that mediate approach behavior (Schroll et al. 2006; Mao and Davis 2009; Liu et al. 2012). Therefore, MBONs and DANs provide an excellent model for studying subcellular-specific targeting of dendrites and axons. However, the mechanisms that guide the projection of MBON dendrites and DAN axons to specific compartments remains unclear.

**Fig. 1.**
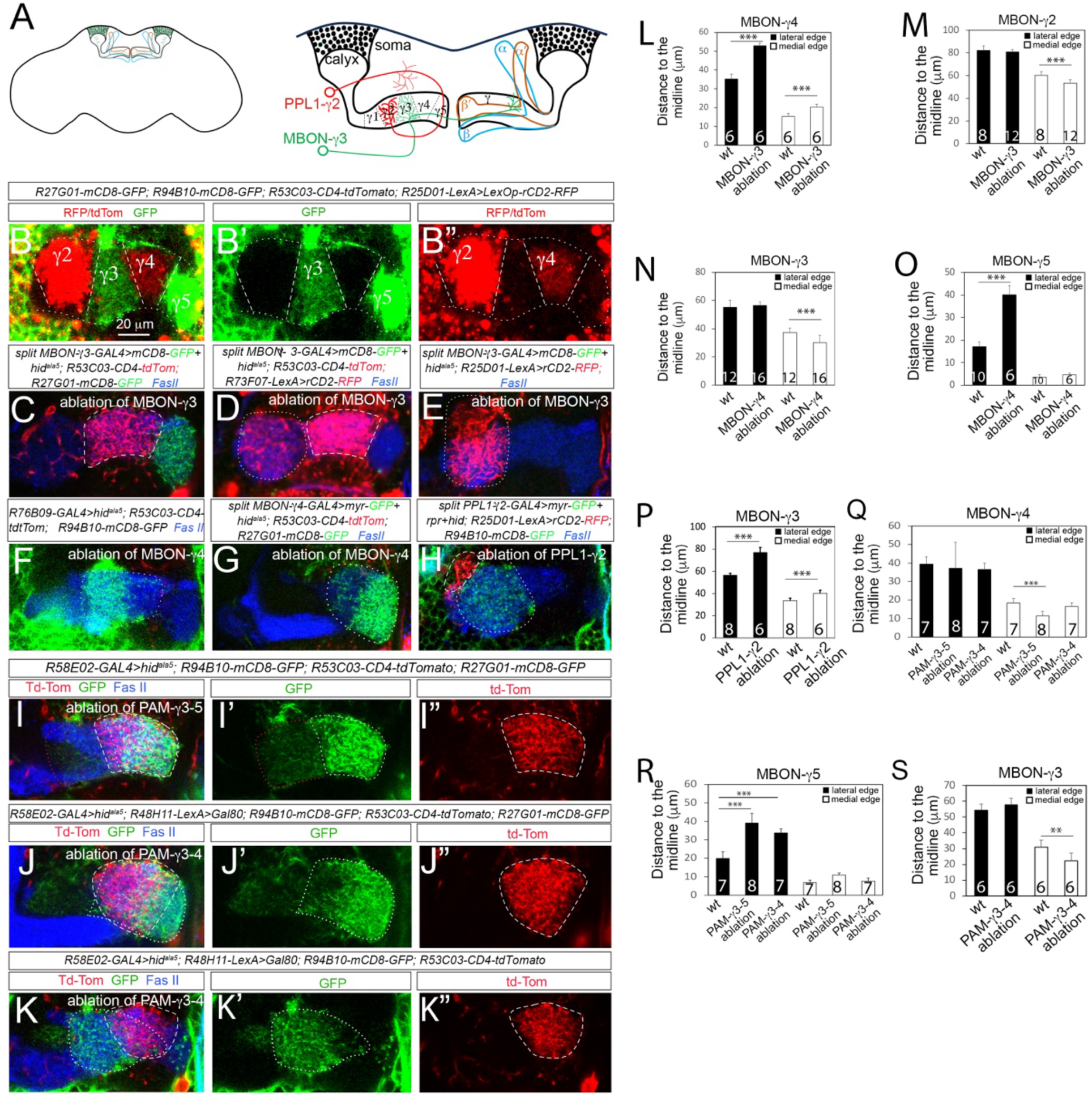
Ablation of MBONs or DANs leads to expansion of the dendritic fields of MBONs. (A) Schematic diagrams of the mushroom body (MB) in the fly brain (left) and an enlarged view of the MB that shows five compartments in the γ lobe and targeting of MBON-γ3 dendrites to the γ3 compartment and PPL1-γ2 axons to the γ2 compartment (right). (B-B”) MBON dendrites in the γ2 through γ5 compartments labeled with alternative fluorescent colors. MBON-γ1 dendrites are not shown in this focal plane. Dotted lines outline the γ lobe and dashed lines demarcate the boundaries between neighboring compartments. (C-E) Ablation of MBON-γ3 neurons by Hid^ala5^ leads to expansion of MBON-γ4 dendrites (stronger RFP, dashed circles) into the γ3 compartment (C), reaching the medial edge of PPL1-γ2 axons (weaker RFP, dotted circle) (D), and slight medial expansion of MBON-γ2 dendrites (dotted circle) (E). (F-G) Ablation of MBON-γ4 neurons results MBON-γ3 dendrites (dotted circle) (F) and MBON-γ5 dendrites (dashed circle) (G) projecting into the γ4 compartment. (H) MBON-γ3 dendrites (dotted circle) project into the γ2 compartment when PPL1-γ2 neurons are ablated. MBON-γ2 dendrites (dashed circle) are displaced from the γ2 compartment and do not overlap with MBON-γ3 dendrites. (I-I”) Ablation of PAM-γ3-5 neurons leads to expansion of MBON-γ4 dendrites (dashed circles) and MBON-γ5 dendrites (labeled with stronger GFP, white dotted circles) into each other’s territories. MBON-γ3 dendrites are labeled with weaker GFP (dotted circles in magenta). (J-K”) Ablation of PAM-γ3-4 alone results in projection of MBON-γ5 dendrites (dotted circles) (J-J”) and MBON-γ3 dendrites (dotted circles) (K-K”) into the γ4 compartment and overlap with MBON-γ4 dendrites (dashed circles). (L-S) Quantifications of the distance from the lateral (filled bars) or medial (open bars) edges to MBON dendritic fields to the midline of the brains in brains with indicated genotypes. Numbers on individual bars represent the number of brain lobes. ***, *p*<0.001; **, *p*<0.01; compared to the wild type; student *t*-test.

Here, we investigated if repulsion between neighboring compartments was required to restrict the projection of MBON dendrites and DAN axons to their specific compartments and tested the role of Slit in mediating the repulsion. We further examined the functional consequences of defects of compartment-specific targeting of MBON dendrites.

## RESULTS

### Compartment-specific targeting of MBON dendrites involves axon- and dendrite-mediated repulsion between neighboring compartments

To investigate mechanisms regulating the compartment-specific targeting of MBON dendrites, we focused on those projecting their dendrites to the MB γ lobe in this study. we first generated transgenic lines, *R27G01-mCD8-GFP*, *R94B10-mCD8-GFP*, *R93D10-mCD8-GFP, and R53C03-CD4-tdTom* to labele MBON-γ5β’2a (MBON-γ5), MBON-γ3β’1 (MBON-γ3), MBON-γ1pedc>α/β (MBON-γ1), and MBON-γ4>γ1,γ2 (MBON-γ4) neurons, respectively (Supplemental Fig. S1A-C’, E-E’). *R27G01*, *R94B10*, *R93D10,* and *R53C03* are enhancer fragments that are active in their respective MBONs (Jenett et al. 2012). In addition, we used the expression of *LexAop-rCD2-RFP* driven by *R25D01-LexA* to label MBON-γ2α’1 (MBON-γ2) neurons (Supplemental Fig. S1D-D’). By combining all these transgenes, we could simultaneously label MBON dendrites targeted to all the five γ lobe compartments with alternate fluorescent colors (Fig. 1B-B”), enabling easy detection of targeting defects of MBON dendrites. Furthermore, these transgenic lines allow gene manipulation in specific neurons using the GAL4-UAS system without interfering the labeling of MBON dendrites.

To test if repulsion between neighboring compartments governs compartment-specific targeting of MBON dendrites, we systematically ablated MBONs or DANs projecting their dendrites or axons to the γ lobe by expressing *UAS-rpr* (*reaper)* + *UAS-hid* (*head involution defective*, *hid*) or *UAS-hid^Ala5^* alone. Meanwhile, we labeled MBONs or DANs using our transgenic lines or existing LexA lines for examining their targeting. We focused on MBON dendrites in compartments γ2 through γ5 for our analyses. We quantified defects of compartment-specific targeting of MBON dendrites by measuring the distance from their lateral or medial edge of their dendritic field to the midline of the brain. Our ablation results show that:

1. Ablation of MBON-γ3 neurons led to dramatic expansion of MBON-γ4 dendrites into the g3 compartment, reaching the lateral edge of the PPL1-γ2α’1 (PPL1-γ2) axonal field (Fig. 1C-D, L). Additionally, MBON-γ2 dendrites also expanded slightly toward the midline but significantly (Fig. 1E, M).
2. Ablation of MBON-γ4 neurons caused MBON-γ3 and MBON-γ5 dendrites to project into the γ4 compartment (Fig. 1F-G, N-O).
3. Ablation of MBON-γ2 or MBON-γ5 neurons did not affect the targeting of MBON-γ3 dendrites or MBON-γ4 dendrites (Supplemental Fig. S2A-D).
4. Ablation of PPL1-γ2 neurons led to projection of MBON-γ3 dendrites into the γ2 compartment and displacement of MBON-γ2 dendrites from the γ2 compartment without overlapping with each other (Fig. 1H, P).
5. Simultaneous ablation of PAM-γ3, PAM-γ4, and PAM-γ5 DANs led to dramatic expansion of MBON-γ4 and MBON-γ5 dendrites into each other’s territories, resulting nearly complete overlap between them and loss of the gap between MBON-γ5 dendrites and MBON-γ3 dendrites (Fig. 1I-I”, Q-R, Supplemental Fig. S2G-H’).
6. Ablation of PAM-γ3 and PAM-γ4 neurons only by simultaneously expressing GAL80 to inhibit the expression of *UAS-hid^ala5^* in PAM-γ5 neurons led to projection of MBON-γ5 and MBON-γ3 dendrites into the γ4 compartment, but did not affect MBON-γ4 dendrite targeting (Fig. 1J-K”, Q-S).
7. Ablation of PAM-γ3 neurons alone did not affect the targeting of MBON-γ3 or MBON-γ4 dendrites (Supplemental Fig. S2E-F).

Together, these ablation results demonstrate that restricting MBON dendrites to their specific compartments critically relies on repulsions between neighboring compartments. The repulsion can be mediated by either MBONs or DANs. These neurons include MBON-γ3, MBON-γ4, PPL1-γ2, PAM-γ4, and PAM-γ5 neurons that can repel MBON dendrites from neighboring compartments. However, since MBON dendrites still projected to the γ lobe when they expanded into neighboring compartments, the γ lobe likely provides attractive cues to guide MBON dendrites to the γ lobe.

### PPL1-γ2 and PAM-γ5 neurons express Slit to repel MBON dendrites and DAN axons from the γ3 and γ4 compartments

What molecules could mediate the repulsion between MBON dendrites or/and DAN axons from two neighboring compartments? Interestingly, previous studies showed that a GAL4 enhancer trap line, *slit-*GAL4^NP2755^ (Tanaka et al. 2008) is expressed in PPL1-γ2 and PAM-γ5 neurons (Fig. 2A-A”). In addition, a GAL4 line driven by a *slit* enhancer fragment *R32F01* (*R32F01-GAL4*) is also expressed in PPL1-γ2 (Fig. 2B-B”) (Jenett et al. 2012). To confirm the expression of Slit in PPL1-γ2 and PMA-γ5 neurons, we performed immunofluorescent staining using the published Slit antibody (Rothberg et al. 1990). Consistently, we detected Slit protein in the soma of PPL1-γ2 neurons by immunostaining and the staining signal was abolished by RNAi knockdown (Supplementary Figure S3B-B’, D-D’) However, we were not able to determine if Slit is expressed in the γ2 or γ5 compartment by immunostaining γ due to ubiquitous Slit staining signals in the entire MB axonal lobes and the dendritic area of the MB, the calyx (Supplementary Figure S3A-A’, B-B’) as reported previously (Oliva et al. 2016). We did not observe obvious reduction of the staining signal in the MB when *elav-GAL4* drove the expression of *UAS-Slit RNAi* pan-neuronally even though knockdown of Slit with *elav-GAL4* led to targeting defects of MB axonal lobes as previously reported (Supplementary Figure S3E-F) (Oliva et al. 2016). Furthermore, expression of *UAS-Slit RNAi* driven by MB-specific *GAL4-OK107* led to neither reduction of Slit staining nor obvious MB axon targeting defects (Supplementary Figure S3G-H), suggesting that the axon targeting defects of MB neurons resulting from the expression of *UAS-slit RNAi* driven by *elav-GAL4* is likely non-cell autonomous. Finally, expression of *UAS-Slit* driven by MB γ neuron-specific *GAL4^NP21^* could not rescue the targeting defects of MB γ lobe in *slit* mutants (Supplementary Figure S3I-K). Therefore, we conclude that the staining signal in the MB is non-specific, which is consistent with the lack of expression of *slit-GAL4^NP2755^* and *R32F01-GAL4* in the MB neurons.

**Fig. 2.**
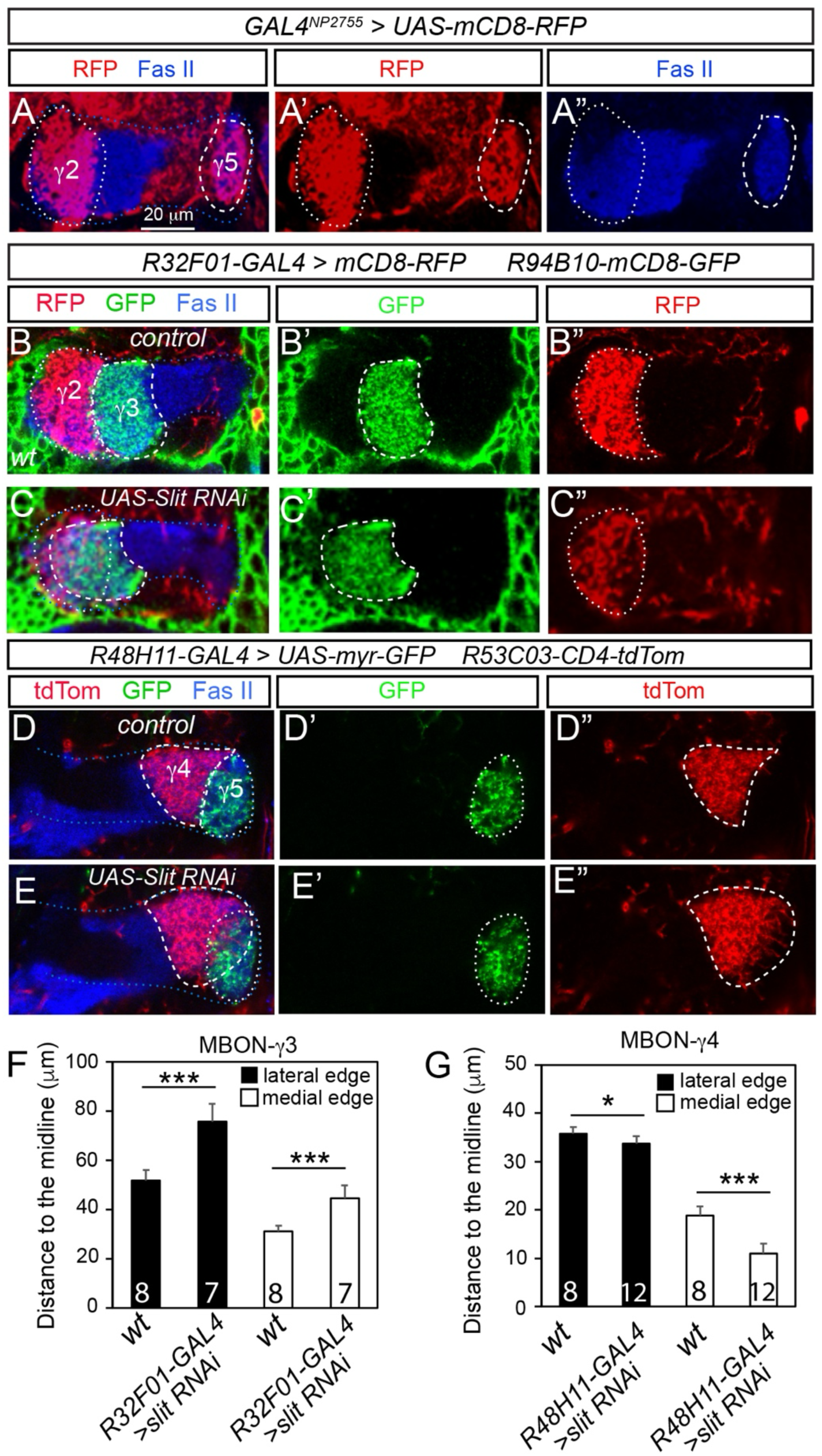
Slit is expressed in PPL1-γ2 and PAM-γ5 neurons to repel MBON-γ3 and MBON-γ4 dendrites. (A-A”) *GAL4^NP2755^* is expressed in both PPL1-γ2 and PAM-γ5 neurons. Their axons in the γ2 and γ5 compartments are outlined with dotted and dashed circles, respectively. (B-C”) Targeting of MBON-γ3 dendrites (GFP, dashed circles) and PPL1-γ2 axons (RFP, dotted circles) in a wild type brain (B-B”) and a brain with Slit RNAi in PPL1-γ2 (C-C”). (D-E”) Projection of MBON-γ4 dendrites (RFP, dashed circles) and PAM-γ5 axons (GFP, dotted circles) in a wild type brain (D-D”) and a brain with Slit RNAi in PAM-γ5 neurons (E-E”). In all images, brains are stained with Fas II antibodies to labeled the γ lobe (outlined with blue dotted lines). (F-G) Quantifications of the distance from the lateral (filled bars) or medial (open bars) edges of the dendritic field of MBON-γ3 (F) and MBON-γ4 (G) neurons to the midline in wild type or Slit knockdown brains. Numbers on individual bars represent the number of brain lobes. ***, *p*<0.001; *, *p*<0.05, compared to control groups; student *t*-test.

To functionally validate if PPL1-γ2 and PAM-γ5 neurons indeed express Slit to repel MBON dendrites from the γ3 and γ4 compartments, we examined how targeting of MBON dendrites and DAN axons in the γ3 and γ4 compartment would be affected when Slit is knocked down in PPL1-γ2 or PAM-γ5 neurons or in *slit* mutant animals. Our results showed that Slit knockdown in PPL1-γ2 caused of MBON-γ3 dendrites to projection into the γ2 compartment, overlapping with PPL1-γ2 neuron axons (Fig. 2B-C”, F). Similarly, Slit knockdown in PAM-γ5 neurons led MBON-γ4 dendrites to project into the γ5 compartment, overlapping with PAM-γ5 axons (Fig. 2D-E”, G). Consistent with Slit knockdown, in *slit^2^/slit^Dui^* mutant brains, MBON-γ3 dendrites also overlapped with MBON-γ2 dendrites and PPL1-γ2 axons (Supplemental Fig. S4A-A”, C-C”). The overlap with PPL1-γ2 axons could rescued by expressing *UAS-slit* in PPL1-γ2 neurons, but expression of *UAS-slit* in MBON-γ2 did not rescue the overlap with MBON-γ2 dendrites (Supplemental Fig. S4B-B”, D-D”), supporting that PPL1-γ2 neurons express Slit to repel MBON-γ3 dendrites. Similarly, MBON-γ4 dendrites overlapped with MBON-γ5 dendrites in *slit^2^/slit^Dui^* mutants (Supplemental Fig. S4E-E”). These findings confirm that PPL1-γ2 and PAM-γ5 neurons express Slit to repel MBON dendrites and DAN axons in the γ3 and γ4 compartments.

### Expression of Robo receptors in MBONs and DANs

Slit functions as a repulsive molecule by binding to its Robo receptors. In *Drosophila*, there are three Robo receptors, Robo1, Robo2, and Robo3 (Kidd et al. 1999; Simpson et al. 2000b). To investigate how Slit repels MBON dendrites in the γ3 and γ4 compartments, we then examined the expression of Robo receptors. We examined Robo1 and Robo3 expression using their existing antibodies, and Robo2 using a Cherry protein trap line Robo2^Cherry^ (Sasse and Klambt 2016). Since the γ lobe undergo remodeling during early pupal stages (Technau and Heisenberg 1982; Lee et al. 1999) and the projection of MBON dendrites observed in adult brains are established during remodeling of the γ lobe at early pupal stages (Truman et al. 2023), we examined their expression at about 2-3 days after puparium formation (APF). We found that all three Robo receptors are expressed in both γ3 and γ4 compartments (Fig. 3A1-A1’, B1-B1’, C1-C1’). By examining the expression of these Robo receptors in the γ3 or γ4 compartments or in their soma after knocking down individual Robo receptors in MBON-γ3, MBON-γ4, PAM-γ3, or PAM-γ4 neurons, we determined the expression of individual Robo receptors in these neurons.

1. Robo1 is expressed in MBON-γ3, MBON-γ4, and PAM-γ4 neurons but not PAM-γ3 as knockdown of Robo1 in MBON-γ3 but not in PAM-γ3 abolished Robo1 in the γ3 compartment, whereas knockdown of Robo1 in MBON-γ4 or PAM-γ4 both reduced Robo1 in the γ4 compartment by 40-50% (Fig. 3A2-A5’, D).
2. Robo2 is expressed in MBON-γ3 and PAM-γ4 neurons but not in PAM-γ3 or MBON-γ4 neurons as Robo2^cherry^ expression is abolished in the γ3 or γ4 compartment when Robo2 was knocked down in MBON-γ3 or PAM-γ4 neurons, respectively (Fig. 2B1-B4’, E). Consistently, Robo2^cherry^ was detected in the soma of MBON-γ3 or PAM-γ4 neurons but abolished by Robo2 RNAi. In contrast, Robo2^cherry^ was not detected in the soma of PAM-γ3 or MBON-γ4 neurons (insets in Fig. 3B1-B6’).
3. Robo3 is expressed in PAM-γ3, MBON-γ3, and PAM-γ4, but not MBON-γ4 neurons as its expression in the γ3 compartment was reduced by about 50% when Robo3 was knocked down in either PAM-γ3 or MBON-γ3 (Fig. 3C1-C3’, F), whereas its expression in γ4 was abolished when Robo3 was knocked down in PAM-γ4 but not MBON-γ4 (Fig. 3C4-C5’, F).

**Fig. 3.**
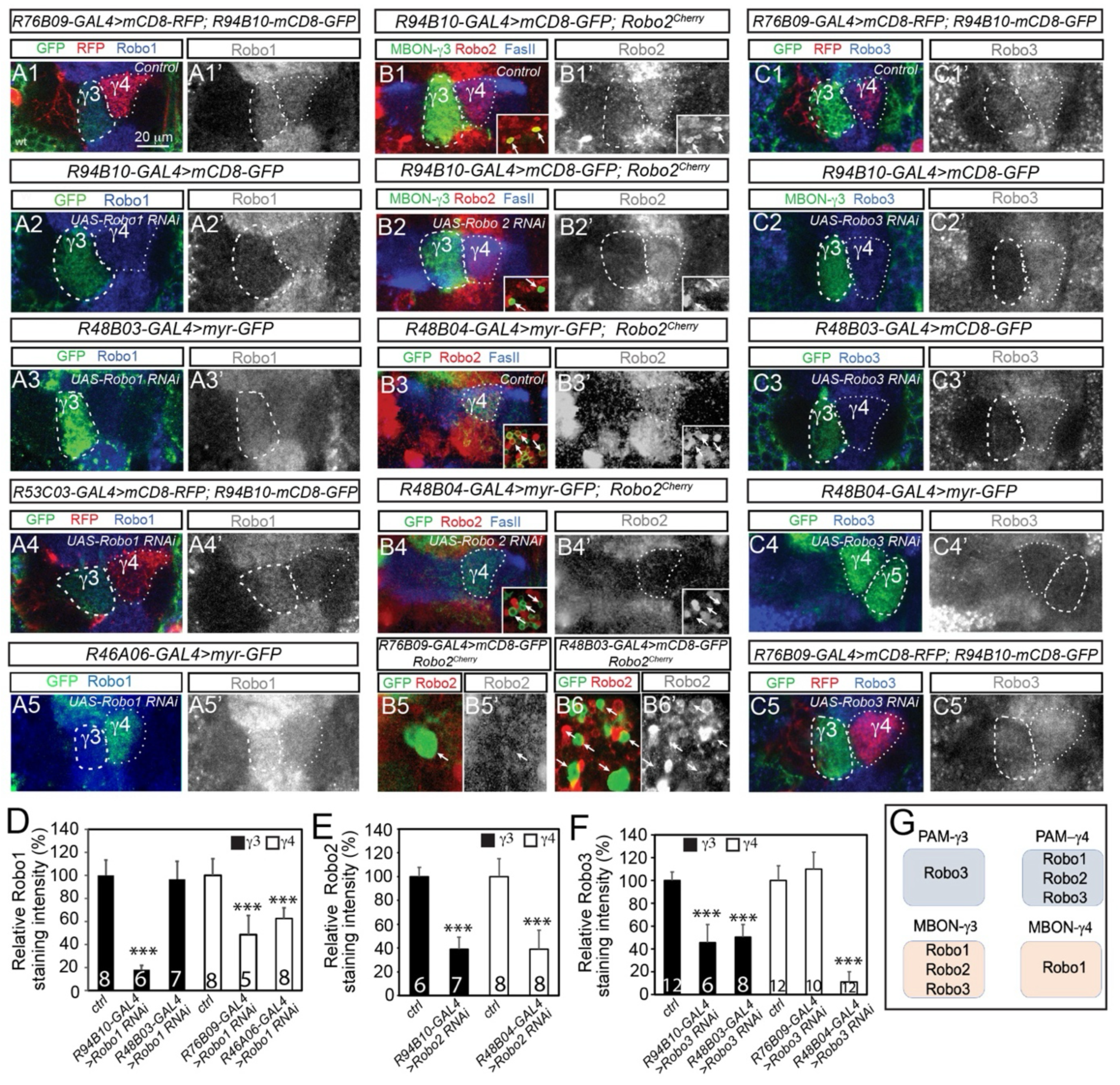
Expression of Robo receptors in MBONs and DANs. In all images, dashed circles outline the γ3 compartment except in (C4-C4’) where they outline the γ5 compartment, and dotted circles the γ4 compartment. (A1-A1’) Endogenous Robo1 proteins are expressed in the γ3 and γ4 compartments in a wild type brain. Dendrites of MBON-γ3 and MBON-γ4 neurons are labeled with GFP and RFP, respectively. (A2-A3’) Expression of Robo1 in the γ3 compartment was abolished by Robo1 RNAi in MBON-γ3 neurons (GFP) using the line BDSC #39027 (A2-A2’) but not in PAM-γ3 neurons (GFP) using the line BDSC #35768 (A3-A3’). (A4-A5’) Robo1 RNAi (BDSC #35768) in MBON-γ4 (RFP) (A4-A4’) or PAM-γ4 (GFP) (A5-A5’) neurons both leads to a significant reduction in Robo1expression in the γ4 compartment. MBON-γ3 neurons are labeled with GFP in (A4). (B1-B1’, B3-B3’) Robo2^Cherry^ is detected in the γ3 and γ4 compartments and in the soma (insets) of MBON-γ3 (GFP in B1-B1’) and PAM-γ4 (GFP in B3-B3’) neurons in wild type brains. (B2-B2’, B4-B4’) Robo2 RNAi in MBON-γ3 (GFP, B2-B2’) or PAM-γ4 (GFP in B4-B4’) neurons dramatically reduced the expression of Robo2^Cherry^ in the γ3 and γ4 compartments, respectively, as well as in their soma (arrows in insets). (B5-B6’) Robo2^Cherry^ is not detected in the soma (arrows) of MBON-γ4 (B5-B5’) or PAM-γ3 (B6-B6’) neurons. (C1-C1’) Expression of Robo3 is detected in both γ3 and γ4 compartments. MBON-γ3 and MBON-γ4 neurons are labeled with GFP and RFP, respectively. (C2-C3’) Robo3 RNAi in MBON-γ3 neurons (GFP (C2-C2’) or PAM-γ3 neurons (GFP) (C3-C3’) leads to a significant reduction in Robo3 expression in the γ3 compartment. (C4-C5’) Robo3 RNAi in PAM-γ4-5 neurons (GFP) (C4-C4’) but not MBON-γ4 neurons (RFP) (C5-C5’) abolishes Robo3 expression in the γ4 compartment. MBON-γ3 neurons are labeled with GFP in (C5). (D-F) Quantifications of relative staining intensities of Robo1 (D), Robo2^Cherry^ (E), and Robo3 (F) in the γ3 and γ4 compartments in the brains with indicated genotypes. Numbers on individual bars represent the number of brain lobes. ***, *p*<0.001, compared to wild type; student *t*-test. (G) A diagram summarizing the expression of Robo1, Robo2, and Robo3 in PAM-γ3, PAM-γ4, MBON-γ3, and MBON-γ4 neurons.

Together, these expression data demonstrate that MBON-γ3 and PAM-γ4 neurons express all three Robo receptors; MBON-γ4 neurons express Robo1 only; and PAM-γ3 neurons express Robo3 only (Fig. 3G).

### Different Robo receptors mediate compartment-specific targeting of MBON dendrites and DAN axons in the γ3 and γ4 compartments

To determine if these Robo receptors indeed regulate compartment-specific targeting of MBON dendrites and/or DAN axons the γ3 and γ4 compartments, we examined how knockdown of these Robo receptors would affect the targeting of their dendrites or axons. Since different Robo receptors could function redundantly (Rajagopalan et al. 2000b; Simpson et al. 2000a), we knocked down different Robo receptors individually or simultaneously. Our results showed that:

1. In MBON-γ3 neurons, Robo1 knockdown with two independent lines (Bloomington *Drosophila* Stock Center (BDSC) lines #35768 and #39027) caused MBON-γ3 dendrites to project into the γ2 compartment, with the line #35768 giving more severe phenotypes as indicated by invasion of more MBON-γ3 dendrites into the γ2 compartment (measured by the relative GFP intensities) and displacement of MBON-γ2 dendrites from the γ2 compartment (Fig. 4A-C, S-T). In contrast, double knockdown of Robo2 and Robo3 did not affect MBON-γ3 dendrite targeting (Fig. 4D, S). However, combining Robo2 or Robo3 knockdown with the weaker Robo1 RNAi line (#39027) produced phenotypes comparable to the stronger Robo1 RNAi line (#35768) (Fig. 4E-F, S-T). Therefore, Robo1 is critical for restricting MBON-γ3 dendrites to the γ3 compartment, but Robo2 and Robo3 provide partial redundancy.
2. In MBON-γ4 neurons, knockdown of Robo1 with either the line #35768 or #39027 caused similar projection of MBON-γ4 dendrites into the γ5 compartment, overlapping with MBON-γ5 dendrites, though most MBON-γ4 dendrites remained in the γ4 compartment (Fig. 4G-H’, U-V). In contrast, double knockdown of Robo2 and Robo3 neither affected the projection of MBON-γ4 dendrites (Fig. 4I) nor enhanced the Robo1 knockdown phenotypes (Fig. 4J-J’, U-V), consistent with no expression of Robo2 or Robo3 in MBON-γ4 neurons. These results suggest that only Robo1 is required to prevent MBON-γ4 dendrites from projecting into the γ5 compartment. However, given that most Robo1 knockdown MBON-γ4 dendrites remained in the γ4 compartment, it is likely that other mechanisms also help restrict MBON-γ4 neuron dendrites to the γ4 compartment.
3. In PAM-γ3 and PAM-γ4 neurons, knockdown of Robo3 caused their axons to project into the γ2 and γ5 compartments, respectively, although most of their axons remained in the γ3 or γ4 compartment (Fig. 4K-L, Q-R’, W-X). In PAM-γ4 neurons, the Robo3 knockdown phenotypes were further enhanced by simultaneous knockdown of Robo1 or/and Robo2 receptors although double knockdown of Robo1 and Robo2 had no obvious phenotypes (Fig. 4M-P, X-Y). Therefore, PAM-γ3 and PAM-γ4 neurons rely primarily on Robo3 to restrict their axons to their specific compartments, but Robo1 and Robo2 have partial redundant roles in PAM-γ4 neurons.

**Fig. 4.**
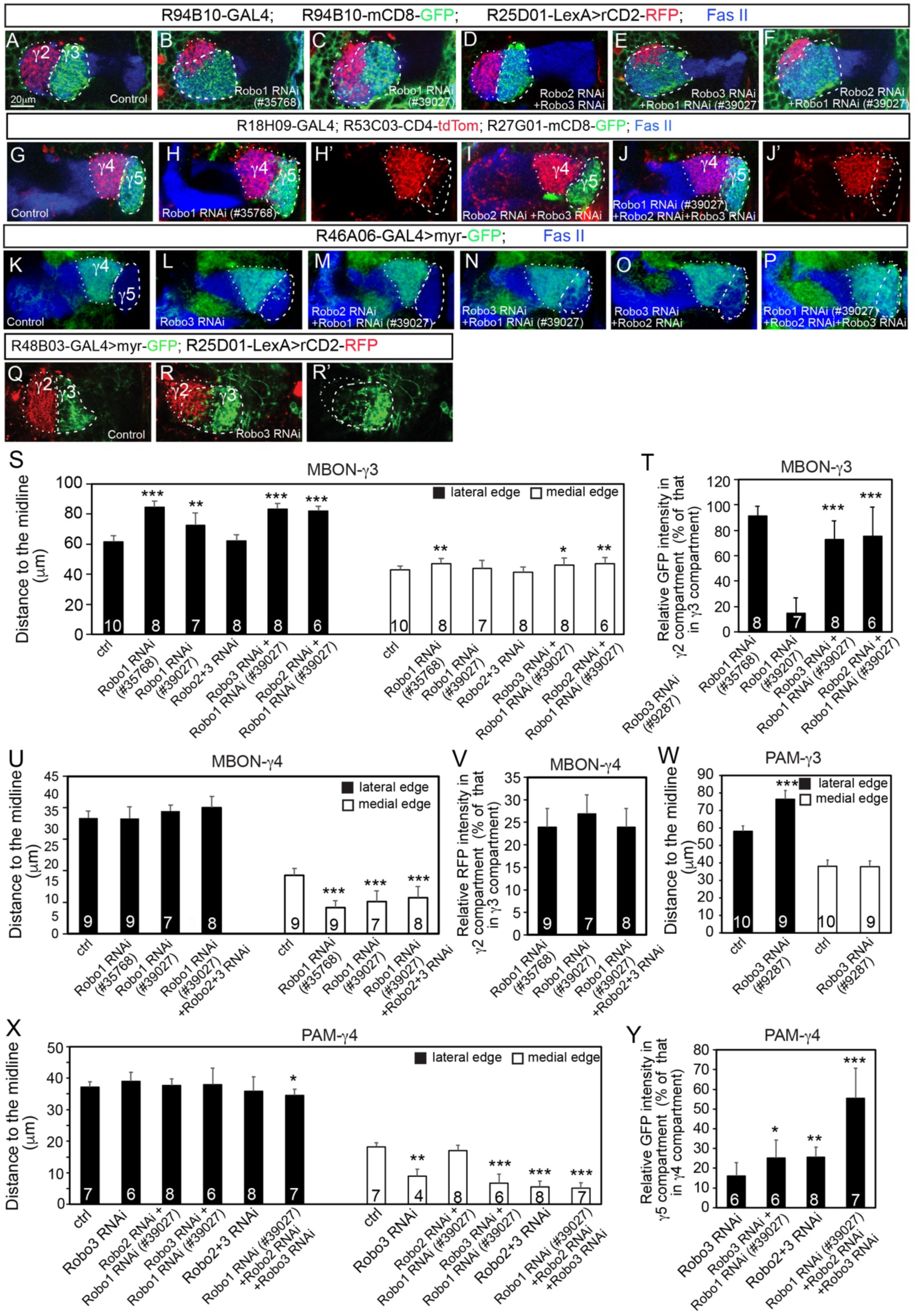
Different Robo receptors are required to prevent the projection of MBON-γ3 and MBON-γ4 dendrites and of PAM-γ3 and PAM-γ4 axons into neighboring compartments. (A-F) Projection of MBON-γ3 dendrites (GFP, dashed circle) and MBON-γ2 dendrites (RFP, dotted circle) in a wild type brain (A), or brains with Robo1 knockdown (with two independent lines) (B-C), double knockdown of Robo2 and Robo3 (D), Robo1 and Robo3 (E), or Robo1 and Robo2 (F) in MBON-γ3 neurons. (G-J’) Projection of MBON-γ4 dendrites (RFP, dotted circles) in a wild type brain (G), or brains with Robo 1 knockdown (H-H’), Robo2 and Robo3 double knockdown (I), and Robo1, Robo2, and Robo3 triple knockdown (J-J’) in MBON-γ4 neurons. MBON-γ5 dendrites are labeled with GFP (dashed circles). (K-P) Projection of PAM-γ4 axons (GFP, dotted circles) in a wild type brain (K), or brains with Robo3 knockdown alone (L), double knockdown of Robo1 and Robo2 (M), Robo1 and Robo3 (N), Robo2 and Robo3 (O), or triple knockdown of Robo1, Robo2, and Robo3 (P) in PAM-γ4 neurons. The γ5 compartment is outlined with dashed circles. (Q-R’) Projection of axons (GFP, dashed circles) of wild type (Q) and Robo3 knockdown (R-R’) PAM-γ3 neurons. MBON-γ2 dendrites are labeled with RFP (dotted circles). (S, U, W, X) Quantifications of the distance from the lateral or medial edge of the dendritic/axonal field of MBONs/DANs with knockdown of different Robo receptors to the midline in the brains. Numbers on individual bars represent the number of brain lobes. ***, *p*<0.001; **, *p*<0.01; *, *p*<0.05, compared to wild type; student *t*-test. (T, V, Y) Quantifications of relative GFP/RFP intensities in the indicated compartments in brains with knockdown of different Robo receptors in MBON-γ3 (T), MBON-γ4 (V), or PAM-γ4 neurons. Numbers on individual bars represent the number of brain lobes. ***, *p*<0.001; **, *p*<0.01; *, *p*<0.05, compared to control groups; student *t*-test.

Together, these RNAi knockdown phenotypes demonstrate that Robo receptors are crucial for restricting the projection of MBON-γ3 and MBON-γ4 dendrites and of PAM-γ3 and PAM-γ4 axons to their specific compartments, but different neurons use different Robo receptors for regulating the compartment-specific targeting of their dendrites or axons.

### Slit-mediated repulsion is required only during early pupal stages for establishing, but not maintaining, compartment-specific targeting of MBON-γ3 dendrites

Since the γ lobe undergoes remodeling during early pupal stages (Lee et al. 1999), and MBON dendrites undergo reorganization during metamorphosis at early pupal stages (Truman et al. 2023), we next tried to determine when Slit-mediated repulsion between MBON-γ3 dendrites and PPL1-γ2 axon occurs and whether Slit is continuously required to maintain the compartment-specific targeting of MBON-γ3 dendrites. We examined PPL1-γ2 axon and MBON-γ3 dendrite projections at larval and early pupal stages in wild type and Slit knockdown brains. In wild type 3^rd^ instar larval brains, PPL1-γ2 (or called DAN-d1 in larval brains) axons are projected to the lateral appendix, a short axonal lobe at the junction of the medial and vertical lobes of the MB γ neurons, while MBON-γ3 (or called MBON-h1 in larval brains) dendrites are projected to the middle of the medial lobe of the MB γ neurons, with a big gap between them (Fig. 5A-A”) (Truman et al. 2023). Between 0-14 hrs APF, when axons of larval γ neurons are being pruned, MBON-γ3 dendrites gradually approached PPL1-γ2 axons (Fig. 5B-C). By 16-18 hrs APF when the γ lobe was completely pruned, MBON-γ3 dendrites physically contacted PPL1-γ2 axons (Fig. 5D-E), but they never crossed the boundary to overlap with PPL1-γ2 axons afterwards (Fig. 5F-G), indicating that after they contact with each other, Slit secreted from PPL1-γ2 neurons prevents MBON-γ3 dendrites from projecting further laterally. By 30 hrs APF when the γ lobe had re-extended, MBON-γ3 dendrites and PPL1-γ2 axons had largely established their adult projection patterns (Fig. 5G).

**Fig. 5.**
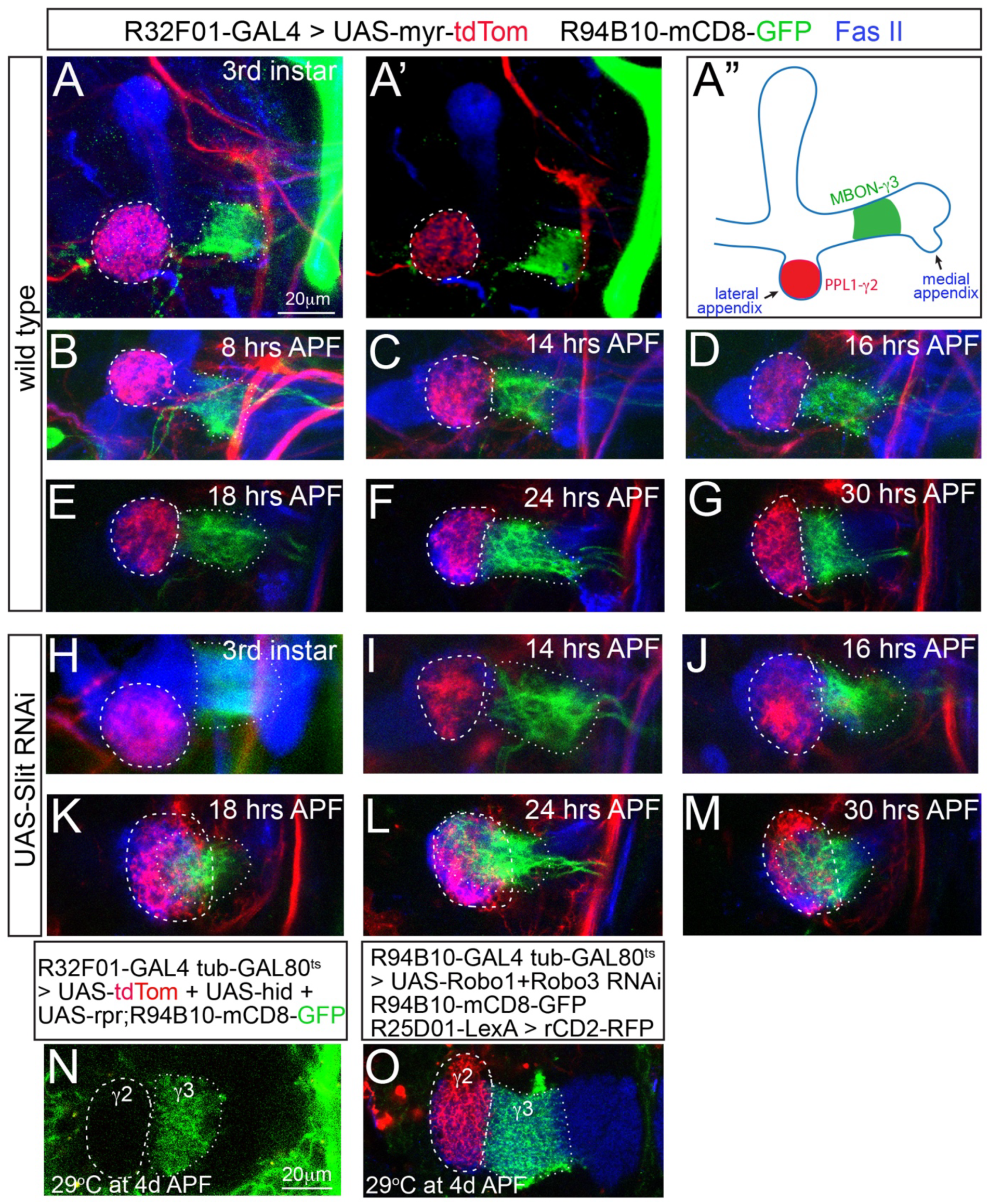
Slit is required at early pupal stages to prevent the projection of MBON-γ3 dendrites into the γ2 compartment. (A-M) Projection of MBON-γ3 dendrites (GFP, dotted circles) and PPL1-γ2 axons (RFP, dashed circles) at the 3^rd^ instar larval stage (A-A”, H) and early pupal stages (B-G, I-M) in wild type brains (A-G) or in brains with Slit RNAi in PPL1-γ2 neurons (H-M). (A’) is a 3D-rendering of the same image in (A) to better show the gap between PPL1-γ2 axons and MBON-γ3 dendrites. (A”) A schematic drawing of the localizations of PPL1-γ2 axons and MBON-γ3 dendrites in larval MB axonal lobes. (N-O) Ablation of PPL1-γ2 neurons or double knockdown of Robo1 and Robo3 after 4 days APF do not affect the targeting of MBON-γ3 dendrites (GFP, dotted circles) and MBON-γ2 dendrites (RFP, dashed circle in (O)) at 18 days after eclosion. In all images, brains are stained with Fas II antibodies to mark MB axonal lobes.

When Slit was knocked down in PPL1-γ2 neurons, projections of MBON-γ3 dendrite and PPL1-γ2 axons did not show obvious targeting defects from 3^rd^ instar larval stage to 16 hrs APF (Fig. 5H-J). However, from 18 hrs APF onward, MBON-γ3 dendrites extended into the γ2 compartment, increasingly overlapping with PPL1-γ2 axons (Fig. 5K-M), indicating that repulsion of MBON-γ3 dendrites by Slit secreted from PPL1-γ2 neurons occurs at early pupal stages and is contact dependent.

To determine whether Slit is required to maintain the compartment-specific targeting of MBON-γ3 dendrites after establishment, we ablated PPL1-γ2 neurons or simultaneously knocked down Robo1 and Robo3 in MBON-γ3 neurons after 4 days APF using *tub-GAL80^ts^* to temporarily control the expression of *UAS-hid* and *UAS-rpr*, or *UAS-RNAi* transgenes. However, even at 18 days after eclosion, MBON-γ3 dendrites showed no targeting defects (Fig. 5N-O), suggesting that Slit-mediated repulsion is only needed for establishing, but not maintaining, the compartment-specific targeting of MBON-γ3 dendrites.

### The expansion of MBON-γ3 dendrites leads to formation of ectopic synaptic contacts

How would expansion of MBON dendrites into neighboring compartments affect their neuronal connections and behavioral outputs? Given that knockdown of Robo receptors gave strong phenotypes in MBON-γ3 neurons, we focused on MBON-γ3 neurons for functional analyses. MBON-γ3 neurons normally form synaptic contacts with PAM-γ3 neurons but not PPL1-γ2 neurons, which can be confirmed with the GRASP (GFP Reconstitution Across Synaptic Partners) approach that detects synaptic contacts (Feinberg et al. 2008) (Fig. 6A-B_1_”). However, when all three Robo receptors were knocked down in MBON-γ3 neurons, we detected GRASP signals between MBON-γ3 dendrites and PPL1-γ2 axons, overlapping with the presynaptic marker Bruchpilot tagged with mCherry (Brp-mCherry) (Berger-Muller et al. 2013) expressed in PAM-γ3 neurons (Fig. 6C-C_1_”), This indicates that expanded MBON-γ3 dendrites formed ectopic synaptic contacts with PPL1-γ2 axons.

**Fig. 6.**
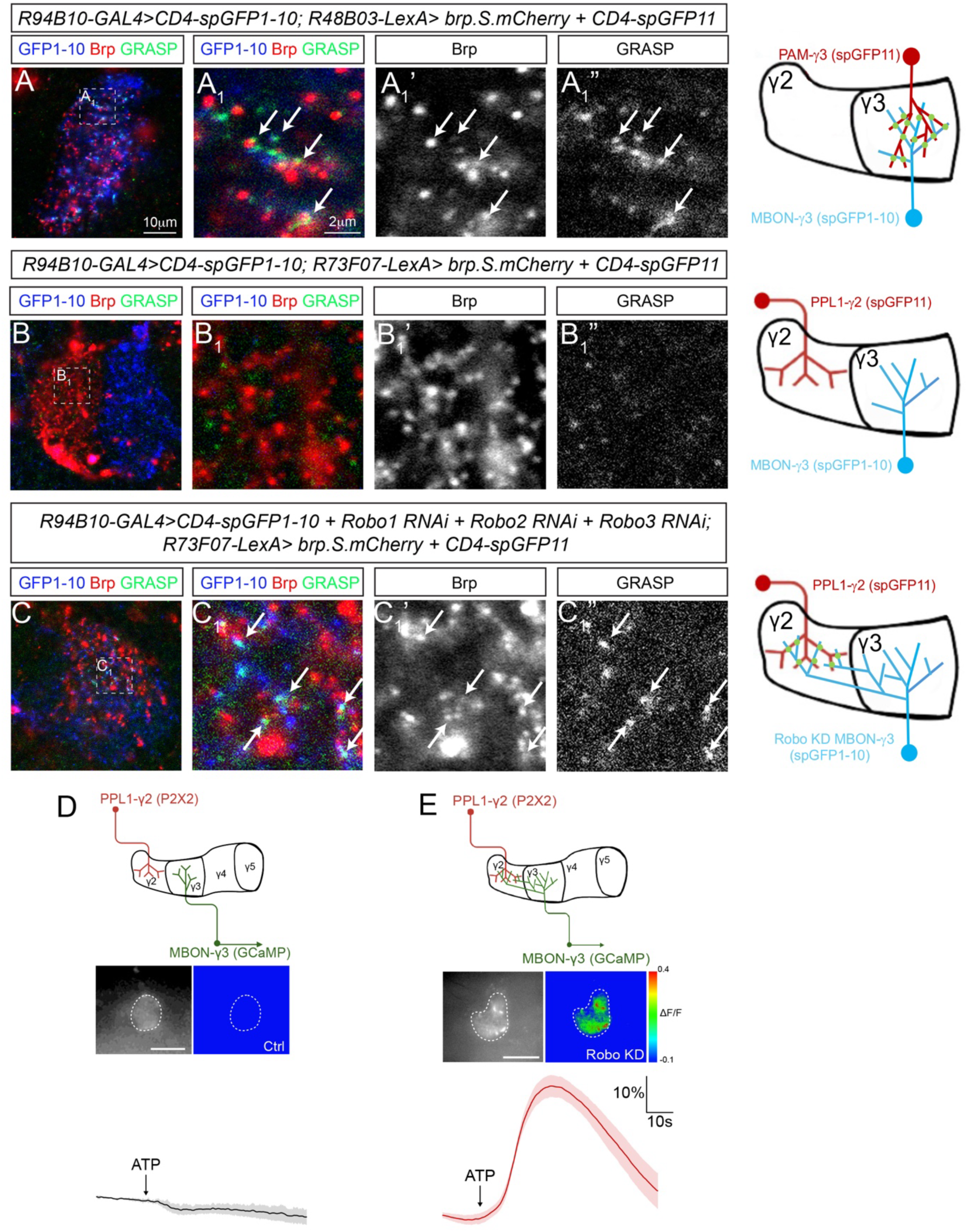
Expansion of MBON-γ3 dendrites leads to formation of ectopic synapses. (A-B_1_”) GRASP signals are detected between MBON-γ3 dendrites and PAM-γ3 axons (A-A_1_”) but not between MBON-γ3 dendrites and PPL1-γ2 axons (B-B_1_”) in wild type brains. (A_1_-A_1_”) and (B_1_-B_1_”) are enlarged views of the area highlighted with dashed squares in (A) and (B), respectively. GRASP puncta (arrows) largely overlap with the presynaptic marker Brp-mCherry expressed in PAM-γ3 in (A_1_-A_1_”). (C-C1”) GRASP signals are detected between MBON-γ3 dendrites and PPL1-γ2 axons when Robo1, Robo2, and Robo3 are simultaneously knocked down in MBON-γ3. (C_1_-C_1_”) is an enlarged view of the area highlighted with a dashed square in (C). GRASP puncta (arrows) largely overlap with the presynaptic marker Brp-mCherry expressed in PPL1-γ2 neurons. (D-E) Activation of P2PX-expressing PPL1-γ2 neurons by ATP elicits strong GCaMP responses in the dendrites of Robo knockdown MBON-γ3 neurons in the presence of TTX (E) but not in wild type MBON-γ3 neurons (D).

To functionally validate the ectopic synaptic contact between MBON-γ3 dendrites and PPL1-γ2 axons, we then performed functional calcium imaging by expressing an ATP-gated ion channel protein P2X2 in PPL1-γ2 neurons and GCaMP8m in MBON-γ3 neurons. We found that activation of P2X2 in PPL1-γ2 neurons by injecting ATP locally into their axonal area in the presence of TTX evoked strong GCaMP8m signals in Robo knockdown but not wild type MBON-γ3 neuron dendrites (Fig. 6D). These results demonstrate that there is direct synaptic transmission from PPL1-γ2 neurons to Robo knockdown MBON-γ3 neurons.

### Knockdown of Robo receptors in MBON-γ3 neurons impacts synaptic plasticity and aversive learning

To assess the functional consequences of the ectopic synaptic connections between Robo knockdown MBON-γ3 neurons and PPL1-γ2 neurons on learning and memory, we monitored odor-evoked activity in MBON-γ3 using GCaMP6f and examined potential defects in synaptic plasticity following aversive olfactory conditioning. We hypothesized that the presence of ectopic synaptic inputs from PPL1-γ2α′1 onto MBON-γ3 in Robo knockdown animals would enhance synaptic plasticity in this compartment.

Using a standard aversive training protocol consisting of a 1-minute odor presentation paired with twelve 1.25-s electric shocks, we observed robust depression of responses to the conditioned odor (CS+) in wild type MBON-γ3 neurons (Supplementary Fig. S5A-B). To our knowledge, this represents the first direct demonstration of synaptic plasticity in MBON-γ3 neurons. Notably, this finding is consistent with previous a report showing decreased synaptic release from Kenyon cells onto MBONs in the γ3 compartment following aversive conditioning, as measured using the acetylcholine reporter GRAB-Ach (Stahl et al. 2022). However, under the same conditions, MBON-γ3 neurons with knockdown of all three Robo receptors did not produce a significant difference in odor-evoked depression compared to control animals (Supplementary Fig. S5C). We reasoned that this lack of effect might reflect a ceiling effect caused by the strength of the training protocol.

To test this possibility, we repeated the experiments using a milder training paradigm consisting of a 20-s odor presentation paired with four 1.25-s electric shocks. Under these conditions, Robo knockdown animals still did not show a significant difference in depression of the CS+ response compared to controls (Fig. 7A-E). However, Robo knockdown resulted in a highly significant potentiation of responses to the CS− odor (the odor not paired with shock; Fig. 7A-E).

**Figure 7.**
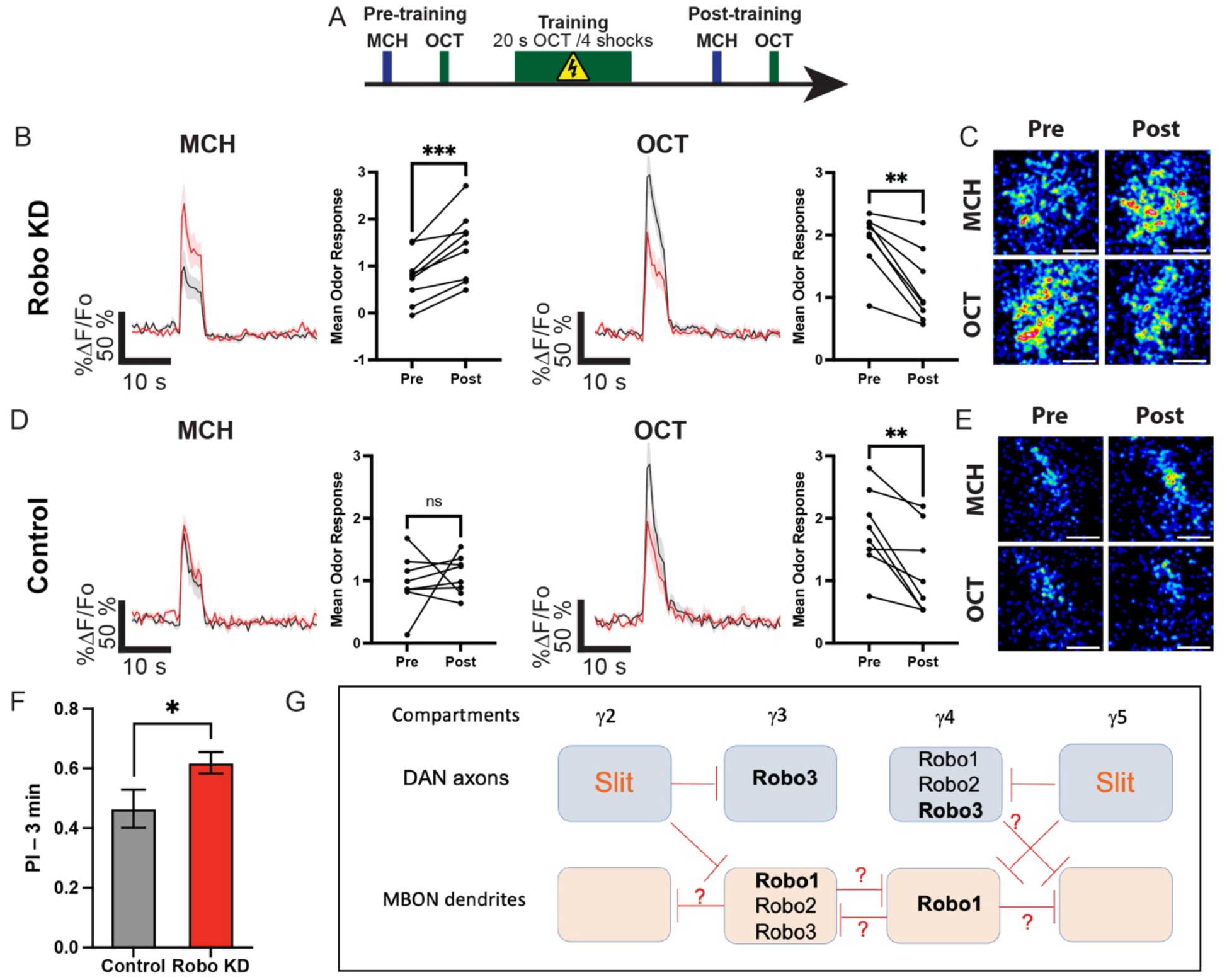
MBON-γ3 neuron memory trace after aversive olfactory conditioning using four electric shocks. (A) Experimental design. Pre-training odor induced calcium response are recorded by presentation of 5-s MCH and OCT with a 30-s ISI; 5 minutes later, flies were aversively trained by pairing 20 s presentation of OCT with 4-90 V shocks. Post-training odor responses were recorded 3 min after. (B, D) GCaMP6f activity in Robo knockdown (B) and control (D) MBON-γ3 neuron dendrites in response to MCH (CS−) (left) and OCT (CS+). (C, E) Pseudocolor images of calcium responses in the MBON-γ3 dendritic areas pre and post-training in Robo knockdown (C) and control animals (E). (F) 3-min memory performance in control and Robo knockdown animals. (G) A schematic diagram of repulsions (red lines) mediated by MBONs or DANs between neighboring compartments. Slit is expressed in PPL1-γ2 and PAM-γ5 neurons. MBONs and DANs in the γ3 and γ4 compartments express different Robo receptors. The Robo receptors in bold play primary roles. In addition to PPL1-γ2 and PAM-γ5 neurons, MBON-γ3, MBON-γ4, and PAM-γ4 neurons also repel MBON dendrites and/or DAN axons in the neighboring compartments.

We next examined whether these physiological changes translated into behavioral consequences by assessing short-term memory performance following the mild training protocol. Consistent with the imaging results, Robo KD animals exhibited a significant enhancement in 3-min memory performance (Fig. 7F). This behavioral effect can be readily explained by the decision-making process during memory retrieval, in which flies choose between the CS+ and CS− odors. Earlier studies have demonstrated that artificial activation of MBON-γ3 drives attraction (Aso et al. 2014b). Therefore, synaptic depression of responses to the CS+ promote avoidance to the punished odor, while potentiation of responses to the CS− increases attraction to the safe odor, together biasing behavior toward avoidance of the conditioned stimulus.

## DISCUSSION

Subcellular specificity is a hallmark of neuronal connections. Using the MBONs and DANs as a model system, we demonstrate here that mutual repulsion between neighboring compartments plays a critical role in regulating compartment-specific targeting of MBON dendrites and DAN axons. We identified Slit as a key repulsive molecule mediating the repulsion between a subset of compartments (Fig. 7G). Failure to restrict the projection of MBON dendrites to their specific compartments due to the loss of Slit-mediated repulsion leads to aberrant synaptic connections and alterations in synaptic plasticity and learning and memory.

### Mutual repulsion between neighboring compartments in compartment-specific targeting of MBON dendrites and DAN axons

Our ablation results reveal that compartment-specific targeting of MBON dendrites critically relies on mutual repulsion between neighboring compartments. The repulsion can be mediated either by MBONs (such as MBON-γ3 and MBON-γ4 neurons) or DANs (such as PPL1-γ2, PAM-γ4, and PAM-γ5 neurons) and can occur between axons, between dendrites, or between axons and dendrites depending on the compartment. Since MBON dendrites and DAN axons are still targeted to the MB axonal lobes when repulsion is impaired, attractive molecules expressed in the MB axonal lobes may guide their targeting to the MB axonal lobes. This attractive mechanism is supported by recent studies showing that in the absence of the α lobe, PPL1-α’2α2 axons fail to target to MB axons (Lin et al. 2022a), whereas disrupting the Dpr12-Dipδ-mediated attraction leads to shortening of the γ lobe and failure of the targeting of PAM-γ4/5 axons to the γ4 and γ5 compartments (Bornstein et al. 2021). However, Dpr12-Dipδ may not be critical for restricting PAM-γ4/5 axons to the γ4 and γ5 compartments as suggested previously (Bornstein et al. 2021) because PAM-γ4 axons could expand into the γ5 compartment when Robo receptors were knocked down. We propose that compartment-specific targeting of MBON dendrites and DAN axons involves a combination of both repulsion and attraction. Attractive molecules expressed in the MB axons first guide the projection of MBON dendrites and/or DAN axons to the appropriate MB axonal lobe. Then the repulsion between neighboring compartments restrict their projection to their specific compartment.

### Slit-Robo mediated repulsion in regulating compartment-specific targeting of MBON dendrites and DAN axons

In this study, we identified Slit as a repulsive molecule that is expressed in PPL1-γ2 and PAM-γ5 neurons to repel both MBON dendrites and DAN axons from the γ3 and γ4 compartments, respectively. The repulsion occurs only during early pupal stages when remodeling/reorganization of the γ lobe and MBON-γ3 dendrites brings MBON-γ3 dendrites physically close to PPL1-γ2 axons and PAM-γ3, PAM-γ4, PAM-γ5, and MBON-γ4 neurons also start to project their axons and dendrites to the γ lobe (Bornstein et al. 2021; Truman et al. 2023). Although Slit is a secreted molecule, Slit-mediated repulsion of MBON dendrites and DAN axons appear to be contact-dependent or very short range, possibly because Slit remains attached to the membrane or immediate adjacent extracellular proteins after secretion as demonstrated previously (Nguyen-Ba-Charvet et al. 2001; Xiao et al. 2011). Our results also show that Slit-mediated repulsion is only required for establishing, not maintaining, the compartment-specific targeting of MBON-γ3 dendrites, It is likely that after establishment, synaptic contacts formed between MBON dendrites, DAN axons, and MB axons may help stabilize their projection.

Interestingly, different MBONs and DANs express different combinations of Robo receptors to mediate Slit repulsion. In *Drosophila* ventral nerve cord and heart, expression of different combinations of Robo receptor determines the strength of the response and the distance of longitudinal axons and different types of heart cells from the midline (Rajagopalan et al. 2000b; Simpson et al. 2000a; Santiago-Martinez et al. 2006). Therefore, It is possible that expression of different Robo receptors in MBONs and DANs may also determine the strength of Slit-mediated repulsion, with the expression of all three Robo receptors conferring the strongest repulsion, and Robo3 or Robo1 alone the weakest. In support of this notion, our results show that knockdown of Robo receptors results in much stronger phenotypes in MBON-γ3 and PAM-γ4 neurons than in MBON-γ4 and PAM-γ3 neurons. Most Robo knockdown MBON-γ4 dendrites and PAM-γ3 axons remain in their original compartment, indicating other mechanisms, such as attraction of PAM-γ3 axons by MBON-γ3 dendrites (our unpublished data), also help restrict their targeting, thus they require weak Slit-mediated repulsion and only need to express Robo1 or Robo3 alone. In contrast, MBON-γ3 dendrites and PAM-γ4 axons may rely solely on Slit-mediated repulsion for compartment-specific targeting and require strong Slit-mediated repulsion, thus they need to express all three Robo receptors. However, different Robo receptors may also have unique roles by activating different downstream signaling pathways as seen in *Drosophila* commissural axons (Rajagopalan et al. 2000a; Simpson et al. 2000b; Spitzweck et al. 2010).

### Functional consequences of the defects of compartment-specific targeting of MBON dendrites

The functional importance of the compartment-specific targeting of MBON dendrites and DAN axons has never been assessed previously due to the lack of tools to perturb the compart-specific targeting. In this study, we show that disrupting repulsions between neighboring compartments leads to expansion of MBON dendrites and DAN axons to neighboring compartments, which allows us to assess the functional significance of the compartment-specific targeting. Using MBON-γ3 neurons as example, we show that expansion of MBON-γ3 dendrites into the γ2 compartment could lead to formation of ectopic synapses, alterations in synaptic plasticity, and aversive learning. The formation of ectopic synapses is evidenced by the ectopic GRASP signals and the direct activation of calcium responses by PPL1-γ2 neurons in Robo knockdown MBON-γ3 dendrites. Such calcium responses activated by DANs (likely through Dop1R2) have also been observed in MBON-α1 and MBON-γ5β′2 dendrites (Sitaraman et al. 2015; Takemura et al. 2017). Thus, MBON dendrites have the capability to form synaptic contacts with DANs in other compartments, but the compartment-specific targeting ensures the specificity of synaptic contacts between MBON dendrites and their partner DAN axons.

Interestingly, knockdown of Robo receptors in MBON-γ3 neurons also enhances the 3-min memory performance. This enhancement seems more contributed by the synaptic potentiation of the response to the CS− odor than by the synaptic depression of the response to the CS+ odor in MBON-γ3 neurons. Although Robo knockdown MBON-γ3 neurons do show synaptic depression of the response to CS+ odor after the aversive training, the depression is comparable to that in wild type MBON-γ3 neurons after both the standard and mild trainings. The reason could be that the neuroplasticity induced by the training happens mostly presynaptically in Kenyon axons as reported previously (Stahl et al. 2022) rather than postsynaptically in MBON-γ3 neuron dendrites. Therefore, although there are ectopic synaptic connections between Robo knockdown MBON-γ3 neurons and PPL1-γ2 neurons, these ectopic synaptic connections may not significantly contributes to the depression in the response to the CS+ odor, but rather may only contribute to the potentiation of the response to the CS− odor. Similar to MBON-γ3 neurons, optogenetic activation of MBON-γ2 leads to attraction (Aso et al. 2014b). Furthermore, aversive olfactory conditioning also results in robust synaptic depression of MBON-γ2 in response to the trained odor, shifting behavior toward aversive (Berry et al. 2018). In this study, we show that Robo knockdown MBON-γ3 dendrites push MBON-γ2 dendrites but not the PPL1-γ2 axons away from the γ2 compartment, which could potentially result in reduced synaptic connections between PPL1-γ2 and its partner MBON-γ2 dendrites. However, this reduced connection would not explain the observed memory phenotype, as it would likely impair learning. However, a caveat to this discussion is that we did not examine how the genetic manipulations in MBONs affect the targeting of their axons, which could provide feedback signals to DANs.

## MATERIALS AND METHODS

### Fly husbandry and strains

*UAS-hid^ala5^* (Bergmann et al. 2002) (gift from F. Pignoni) or a combination of *UAS-hid* and *UAS-reaper (rpr)* (Zhou et al. 1997) were used for genetic ablation of MBONs or DANs. LexAop-GAL80 (Pfeiffer et al. 2010) (Bloomington Drosophila Stock Center [BDSC] #32217) was used to inhibit *UAS-hid^ala5^* expression in PAM-γ5 neurons. *UAS-mCD8-GFP* (BDSC #31289), *UAS-myr-GFP* (BDSC #32196), *UAS-mCD8-RFP* (BDSC #32219), *UAS-myr-tdTomato* (BDSC #32221). *LexAop-rCD2-RFP*; *UAS-CD4-spGFP1-10*; *LexAop-CD4-spGFP11* (BDSC #58755) were used for GRASP analyses. *LexAop-brp.s.mCherry* (a gift from W.B. Grueber) (Berger-Muller et al. 2013) was used for labeling presynaptic boutons. *UAS-slit RNAi* (BDSC #31468), *sli^Dui^* (BDSC #9284) and *sli^2^*(BDSC #3266) were used for Slit loss-of-function analyses. *UAS-sli*.D (BDSC stock #67688) was used for rescuing *sli^2^/sli^Dui^* phenotypes. Robo2^Cherry^ (gift from C. Klambt) (Sasse and Klambt 2016) were used for examining Robo2 expression. *UAS-robo1 RNAi* (BDSC stocks #35768 and #39027), *UAS-robo2 RNAi* (BDSC stock #34589), and *UAS-robo3 RNAi* (BDSC stock #9287) were used for knocking down Robo receptors. LexAop-P2x2 (BDSC #76030) (Clowney et al. 2015) was used for activation of PPL1-γ2 neurons with ATP. *UAS-GCaMP6f* (BDSC #42747) and *UAS-GCaMP8m* (BDSC #605072) were used for *in vivo* and *ex vivo* calcium imaging, respectively. For the genetic ablation and RNAi knockdown, animals were raised at 29°C to enhance the efficiency. For genetical ablation of PPL1-γ2 neurons or knockdown of Robo receptors specifically after 4 days APF, *tub-GAL80^ts^* (BDSC #7108) (McGuire et al. 2003) was used in combination with cell type-specific GAL4 lines and animals were kept at 18°C and shifted to 29°C at 4 days APF. GAL4 and LexA lines used for driving the expression of UAS/LexAop-transgenes are listed in the Supplemental Table S1.

### Construction of plasmids and generation of transgenic lines

To generate *R27G01-mCD8-GFP*, *R94B10-mCD8-GFP*, and *R93D10-mCD8-GFP MBON-γ1* constructs, the enhancer fragments, R27G01, R94B10, and R93D10, were amplified by PCR from genomic DNAs using primers flanked by BglII and KpnI sites, and cloned first into pJet2.1 vector. The enhancer fragments R27G01 and R94B10 were then cut from pJet2.1 with either BglII/KpnI or NotI /NheI and two copies of each were incorporated into the pAPIC-PHsIH-3xmCD8::GFP vector (Han et al. 2011) at BglII/KpnI and NotI/NheI sites, respectively, to replace the *ppk* enhancer fragment. The R93D10 fragment was cut from pJet2.1 with NotI/NheI and a single copy was incorporated into the same vector. The resulting constructs enable these enhancer fragments to drive the expression of three copies of mCD8::GFP.

The *R53C03-CD4-tdTomato* construct was generated using the gateway technology. The R53C03 enhancer fragment was amplified from genomic DNAs by PCR with the incorporation of the attB1 and attB2 sequences at the 5’ and the 3’ ends, respectively. The enhancer fragment was cloned into the donor vector pDONR 211 (Lifetechnology, Carlsbad, CA) at the attP1/attP2 sites through a BP reaction. The generated entry clone was used to transfer the enhancer fragment through a LR reaction to the destination vector pDEST-HemmarR (Han et al. 2011), which allows it to drive CD4-tdTomato expression.

Primers for enhancer fragment amplification were designed based on the sequences available from the Janelia FlyLight website (https://flweb.janelia.org/cgi-bin/flew.cgi) (Jenett et al. 2012) and are listed in Supplemental Table S2. The constructs were injected into *w^1118^ Drosophila* embryos by the Rainbow Transgenic Flies, Inc. (Camarillo, California). A standard P-element transposon protocol was used for generating transformants.

### Fluorescent immunostaining and Confocal microscopy imaging

*Drosophila* larval, pupal, or adult brains were dissected and immunostained using a standard protocol as described (Lee et al. 1999). Primary antibodies and fluorophore-conjugated secondary antibodies used in this study are listed in Supplemental Table S3. Images were collected using a Carl Zeiss LSM780 confocal microscopy and processed with Adobe Photoshop.

### *Ex vivo* calcium imaging of MBON-γ3 neuron dendrites

*Ex vivo* calcium imaging was performed as previously described (Lin et al. 2022b). Adult fly brains were dissected in adult hemolymph-like (AHL) solution (Wang et al. 2003) and immobilized with fine tungsten pins in a Sylgard-based chamber. Prior to imaging, calcium-free AHL was replaced with AHL saline containing 2 mM Ca^2+^ and 1 µM tetrodotoxin (TTX) to block action potential–dependent synaptic transmission and eliminate polysynaptic network activity. To stimulate P2X2 channels in PPL1-γ2 neurons, ATP was locally applied in their axonal area using a fine glass electrode mounted on a micromanipulator and positioned in the target region under visible and fluorescent guidance. 5 mM ATP solution was delivered via precisely controlled pressure pulses using a pressure injector. The pressure required for ATP delivery was empirically determined for each preparation. Images were acquired using an sCMOS camera (Hamamatsu Photonics K.K., Bridgewater, New Jersey) controlled by HCImage Live, at a resolution of 1,024 × 1,024 pixels, a frame rate of 3 Hz, and an exposure time of 100 ms, on an Olympus fluorescence microscope. Calcium dynamics (ΔF/F) were extracted by drawing regions of interest (ROIs). Imaging data were analyzed using ImageJ and plotted using Prism and Igor Pro (WaveMetrics, Lake Oswego, Oregon).

### Aversive olfactory training under microscope and *In vivo* calcium imaging

*In vivo* imaging of calcium responses in MBON-γ3 dendrites following aversive conditioning was carried out as described previously (Cervantes-Sandoval et al. 2017; Martinez-Cervantes et al. 2022). Briefly, a single fly was inserted into a custom-made recording chamber after being aspirated into a metal pipette without anesthesia. Then the eyes were glued to the chamber using myristic acid to fix the head. After the fly was released from the metal pipette, the proboscis was fixed with myristic acid to immobilize the brain. Afterwards, a small square section of dorsal cuticle was removed from the head for imaging, followed by perfusion across the brain with saline (103 mM NaCl, 3 mM KCl, 5 mM HEPES (4-(2-hydroxyethyl)-1-piperazineethanesulfonic acid), 1.5 mM CaCl_2_, MgCl_2_, 26 mM NaHCO_3_, 1 mM NaH_2_PO_4_, 10 mM trehalose, 7 mM sucrose, and 10 mM glucose [pH 7.2]). The GCaMP6f fluorescence response in MBON-γ3 dendrites was detected with one HyD channel (510-550 nm) and imaged with a 20x water-immersion objective and a Leica TCS SP8 II confocal microscope for 2 min at 2 Hz. The odor was delivered starting at 30 sec after imaging initiation.

For preconditioning recordings, the flies were sequentially presented to associated and non-associated odors for 5 sec with 30 sec of clean air in between. 5 min after preconditioning recordings, the flies were trained under microscopy by presentation of a single 1 min (or 20 sec for a milder training paradigm) associated odor pulse with simultaneous 12 (or 4 for a milder training paradigm) 90-V, 1.25 sec electric shocks, followed by 1 min (or 20 sec) non-associated odor without the electric shocks with a 30 sec of clean air between two odors. Post-conditioning odor responses were recorded 5 min after training. Odors were delivered to flies with Teflon tubing after a series of dilution and the electric shocks were delivered with a custom-made shock platform. An Arduino microcontroller (Arduino Uno) was used to control both solenoids that control odor delivery and the Grass stimulator that delivers electric shocks.

### Aversive Olfactory memory test

Groups of 60 flies were first acclimated for ~15 min in a fresh food vial to the environment of a behavioral room dimly lit with red light at 24°C and 65% humidity. They were then transferred into a training tube where they received 30 sec of air, followed by 20 sec of an odor paired with 4 pulses of 90V electric shock (CS+), 30 sec of air, 20 sec of a second odor with no electric shock (CS−), and finally 30 sec of air. For conditioning odors, we bubbled fresh air through 3-octanol (OCT) and 4-methylcyclohexonal (MCH) at concentrations of 0.07% and 0.12% in mineral oil, respectively. To measure three minutes memory, we immediately loaded the flies into a T-maze where they were allowed 2 min to choose between the CS+ odor and the CS−odor. For all experiments, two groups were trained and tested simultaneously. One group was trained with OCT as the conditioned stimulus paired with reinforcer (CS+) and MCH unpaired with reinforcer (CS−), while the other group was trained with MCH as CS+ and OCT as CS−. Each group (60 flies) tested provided a half performance index (half PI): Half PI = ((# flies in CS− arm) – (# flies in CS+ arm)) / (# flies in both arms). A final PI was calculated by averaging the two half PI’s. Since the two groups were trained to opposite CS+/CS− odor pairs, this method balanced out naïve odor biases.

### Quantifications and Statistical analyses

For quantifying the distance from the lateral or medial edges of the dendritic field of MBONs or axonal field of DANs to the midline of the brain, we measured the distance from the midpoint of the medial or lateral edges to the midline using the focal slice in the middle of the z-stack. To quantify Robo expression, the staining intensities of Robo receptors in the γ3 or γ4 compartment in wild type or RNAi knockdown brains were measured using the histogram function in photoshop and normalized by the signal intensity in neighboring γ4 or γ3 compartment, respectively. Student’s *t*-test was used for statistical analyses. Graphs were generated with Microsoft Excel or Prism.

## ACKNOWLEDGEMENTS

We thank Drs. C. Klambt, W.B. Grueber for fly lines; the Bloomington *Drosophila* Stock Center and the TRiP at Harvard Medical School (NIH/NIGMS R01-GM084947), and Vienna *Drosophila* Resource Center for providing RNAi transgenic fly stocks; the Developmental Studies Hybridoma Bank for antibodies; members of the Zhu, Pignoni, and Lin Labs for thoughtful discussion; Rainbow Transgenic Flies Inc for embryo injection; Neuroscience Microscopy Core at Upstate Medical University for providing Zeiss LSM 780 confocal microscopy. This work was supported by the National Institute of Neurological Disorders and Stroke of the National Institutes of Health under Award Number R01NS085232 (S.Z.), R21NS109748 (S.Z.), and National Institute of General Medical Sciences of the National Institutes of Health under the award number of R01GM147917 (I.C.S.).

## AUTHOR CONTRIBUTIONS

Conceptualization, X.D, I.C.S., and S.Z.; Methodology: X.D., P.S., I.C.S., H.H.L. and S.Z.; Investigation: X.D., P.S., I.C.S., and H.H.L.; Data analysis: X.D., P.S., I.C.S., H.H.L., and S.Z.; Writing - Original Draft, S.Z., I.C.S., H.H.L. and Writing - Review & Editing - S.Z., I.C.S., H.H.L., and X.D.; Supervision, S.Z.; Funding, S.Z. and I.C.S.

## DECLARE OF INTERESTS

The authors declare no competing interests.

---

**Supplemental Table S1.** GAL4/LexA lines used in this study.

| Neurons | GAL4/LexA lines | stock # |
| --- | --- | --- |
| MBON-$\gamma 2\alpha'1$ | split MBON-$\gamma 2\alpha'1$-GAL4 | BDSC 68249 |
|  | R25D01-LexA | BDSC 49122 |
| MBON-$\gamma 3$ | R94B10-GAL4 | BDSC 41325 |
| | split MBON-$\gamma 3$-GAL4 | BDSC 68287 |
| MBON-$\gamma 4$ | R18H09-GAL4 | BDSC 48830 |
|  | R53C03-GAL4 | BDSC 38870 |
|  | R76B09-GAL4 | BDSC 46962 |
| | split MBON-$\gamma 4$-GAL4 | BDSC 68309 |
| MBON-$\gamma 5$ | R27G01-GAL4 | BDSC 49233 |
| | split MBON-$\gamma 5$-GAL4 | BDSC 68272 |
| PPL1-$\gamma 2$ | R32F01-GAL4 | BDSC 49359 |
| | split PPL1-$\gamma 2$-GAL4 | BDSC 68279 |
|  |  | BDSC 68290 |
|  | R73F07-LexA | BDSC 52743 |
| PAM-$\gamma 3$ | R48B03-GAL4 | BDSC 50346 |
| | split PAM-$\gamma 3$-GAL4 | BDSC 68251 |
|  | R48B03-LexA | BDSC 54214 |
| PAM-$\gamma 4$ | R46A06-GAL4 | BDSC 50250 |
| PAM-$\gamma 5$ | R48H11-GAL4 | BDSC 50396 |
| | split PAM-$\gamma 5$-GAL4 | BDSC 48316 |
|  | R48H11-LexA | BDSC 52745 |
| PAM-$\gamma 3$-5 | R58E02-GAL4 | BDSC 41347 |
| PAM-$\gamma 4$-5 | R48B04-GAL4 | BDSC 50347 |
| PAM-$\gamma 5$+PPL1-$\gamma 2$ | Gal4-NP2755 | DGRC 113037 |
| pan-neuronal | Elav-GAL4 | BDSC 458 |
| all MB neurons | Gal4-OK107 | BDSC 854 |
| MB $\gamma$ neurons | Gal4-NP21 | DGRC 103496 |

**Supplemental Table S2.**
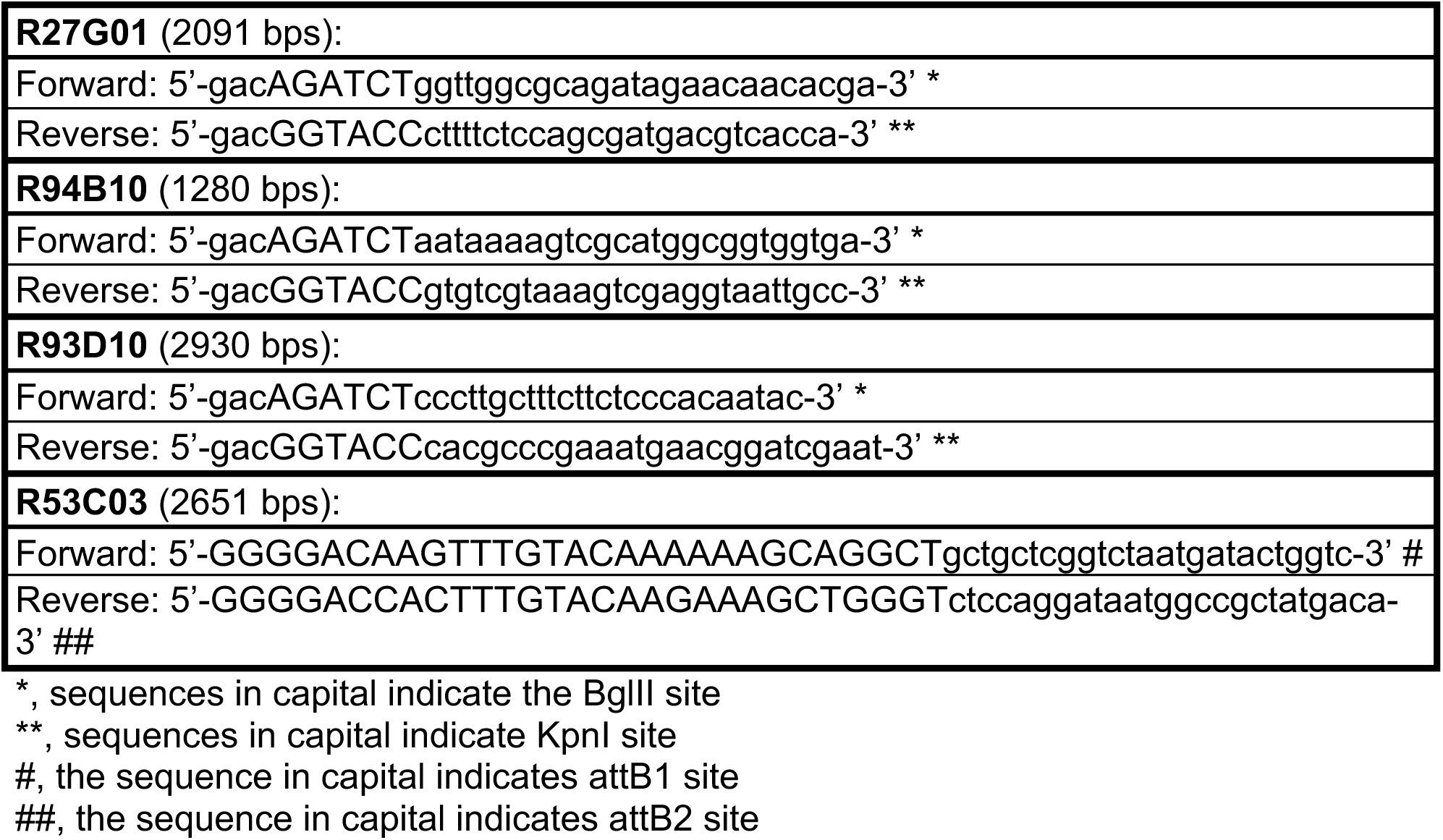
Primers used for amplifying enhancer fragments.

**Supplementary Table S3.**
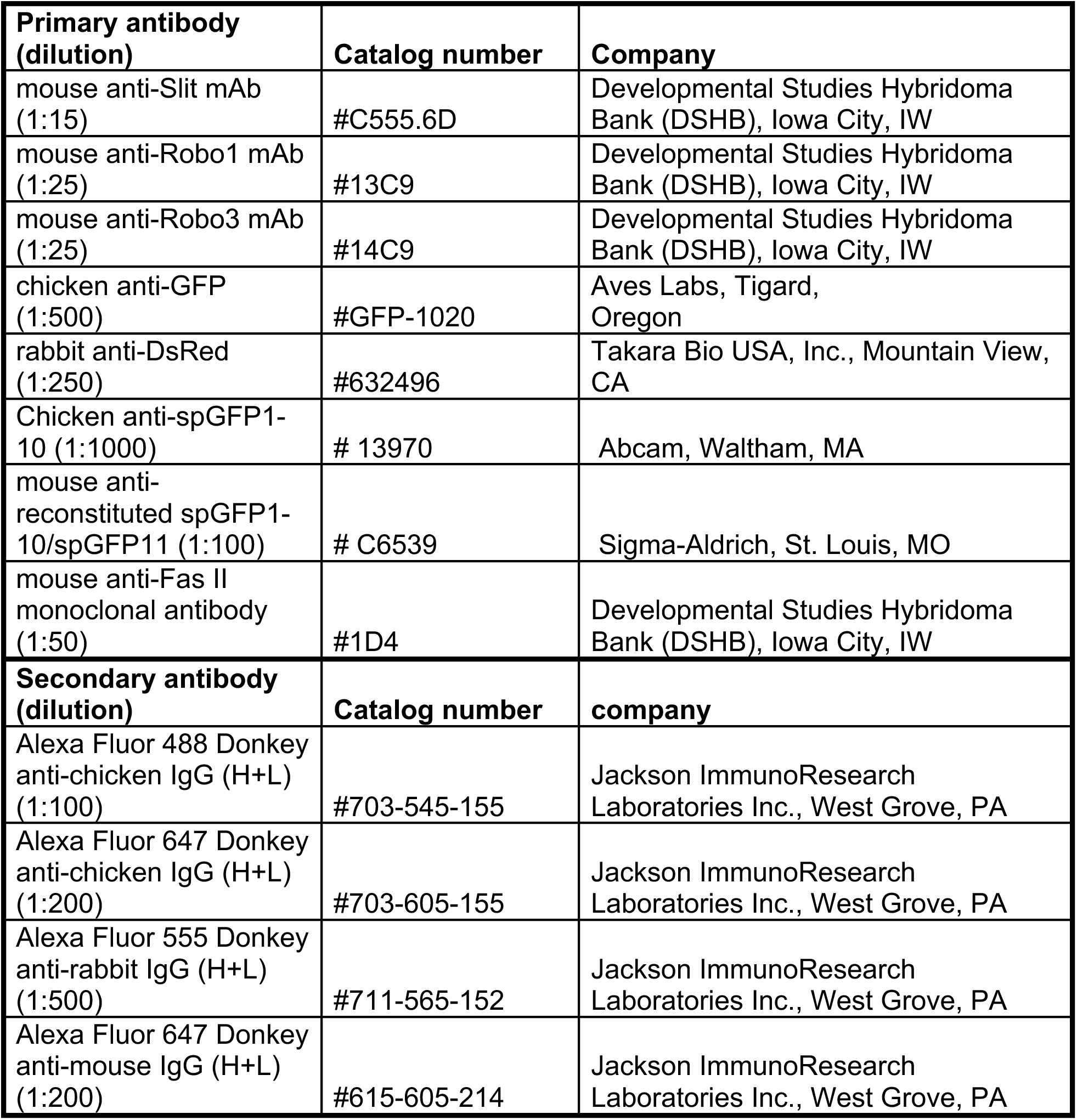
Primary and secondary antibodies used in the study.

**Supplemental Fig. S1.**
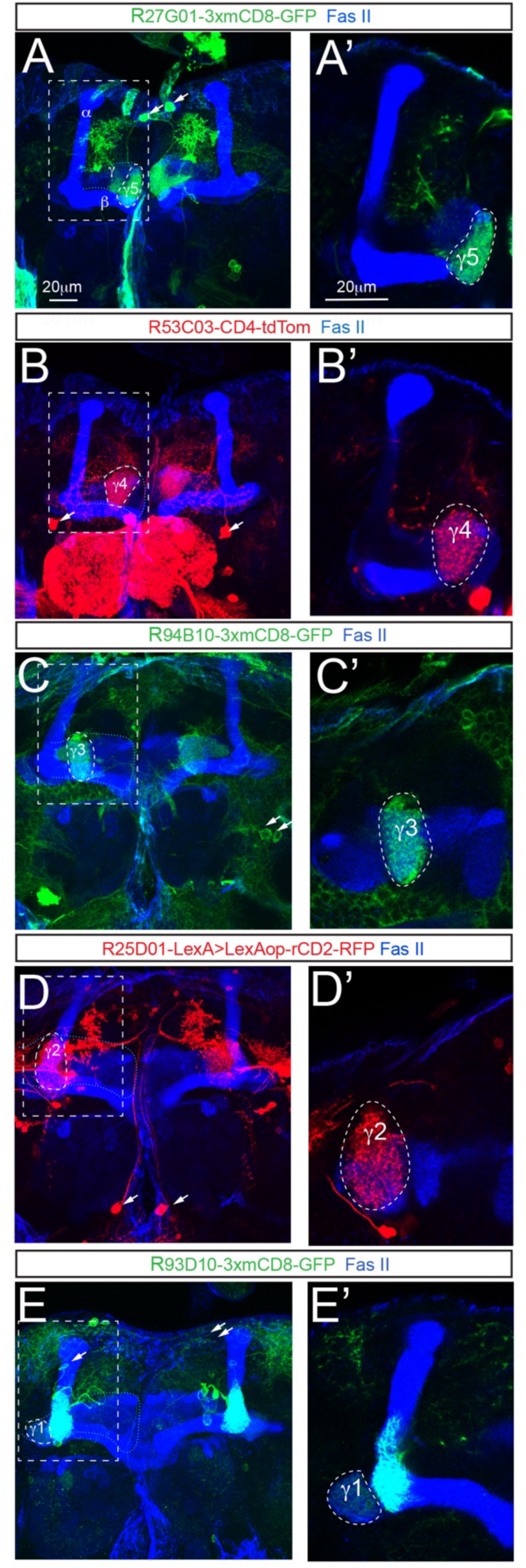
Labeling of different types of MBONs with different fluorescent markers. (A, B, C, D, E) show projected images of MBON-γ5 (A), MBON-γ4 (B), MBON-γ3 (C), MBON-γ2 (D), and MBON-γ1 (E) neurons labeled with R27G01-3xmCD8-GFP, R53C03-CD4-tdTom, R94B10-3xmCD8-GFP, LexAop-rCD2-RFP driven by R25D01-LexA, and R93D10-3xmCD8-GFP, respectively. Brains are counterstained with Fas II antibodies to labeled MB γ and α/β axonal lobes. Arrows point to the soma of MBONs. (A’, B’, C’, D’, E’) show enlarged views of the areas highlighted with dashed rectangles in (A, B, C, D, E) to show dendrites of the corresponding MBONs in their respective compartments in the γ lobe in a single focal slice. Dashed circles outline the dendritic areas of individual MBONs.

**Supplemental Fig. S2.**
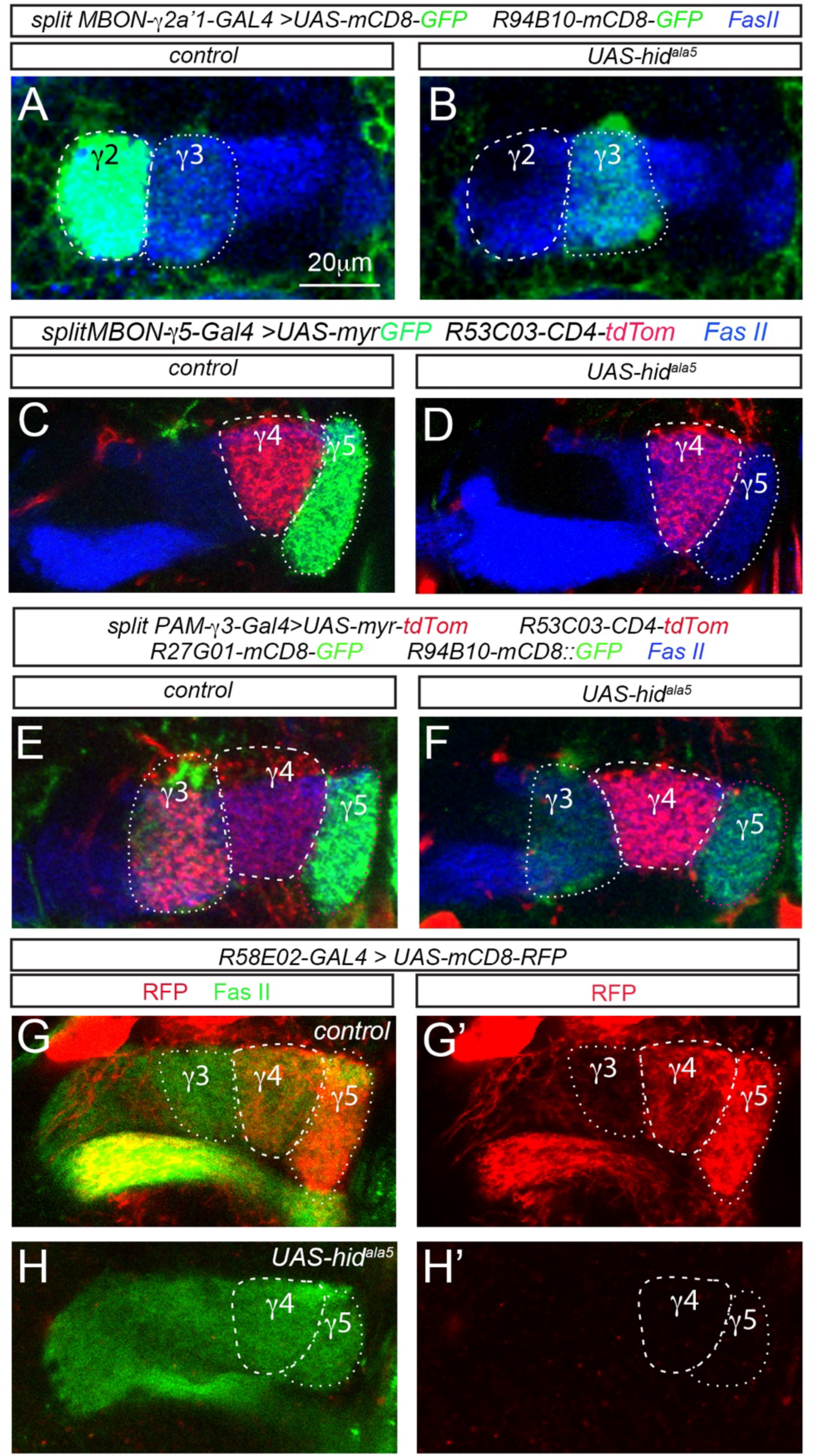
Genetic ablation of MBON-γ2, MBON-γ5, or PAM-γ3 neurons does not affect projection of MBON dendrites in neighboring compartments. (A-B) MBON-γ3 dendrites (weaker GFP, dotted circles) remain in the γ3 compartment as in the wild type brains (A) after ablation of MBON-γ2 neurons (B). Dendrites of MBON-γ2 neurons are labeled with stronger GFP (dashed circles). (C-D) MBON-γ4 dendrites (dashed circles) are similarly targeted to the γ4 compartment in wild type (C) and MBON-γ5-ablated (D) brains. (E-F) Projection of MBON-γ4 dendrites (dashed circle) and MBON-γ3 dendrites (weaker GFP, white dotted circles) remains similar in the wild type (E) and PAM-γ3-ablated (F) brains. PAM-γ3 neurons are labeled with myr-tdTom. MBON-γ5 dendrites are also labeled with stronger GFP in the γ5 compartment (purple circles). (G-G”) mCD8-RFP driven by R58E02-GAL4 labels PAM-γ3-5 axons in the γ3, γ4, and γ5 compartments. (H-H”) Axons of PAM-γ3-5 neurons are absent in the γ lobe after ablation. MB axonal lobes are counterstained with anti-Fas II antibodies in blue (A-F) or green (G-H”).

**Supplemental Fig. S3.**
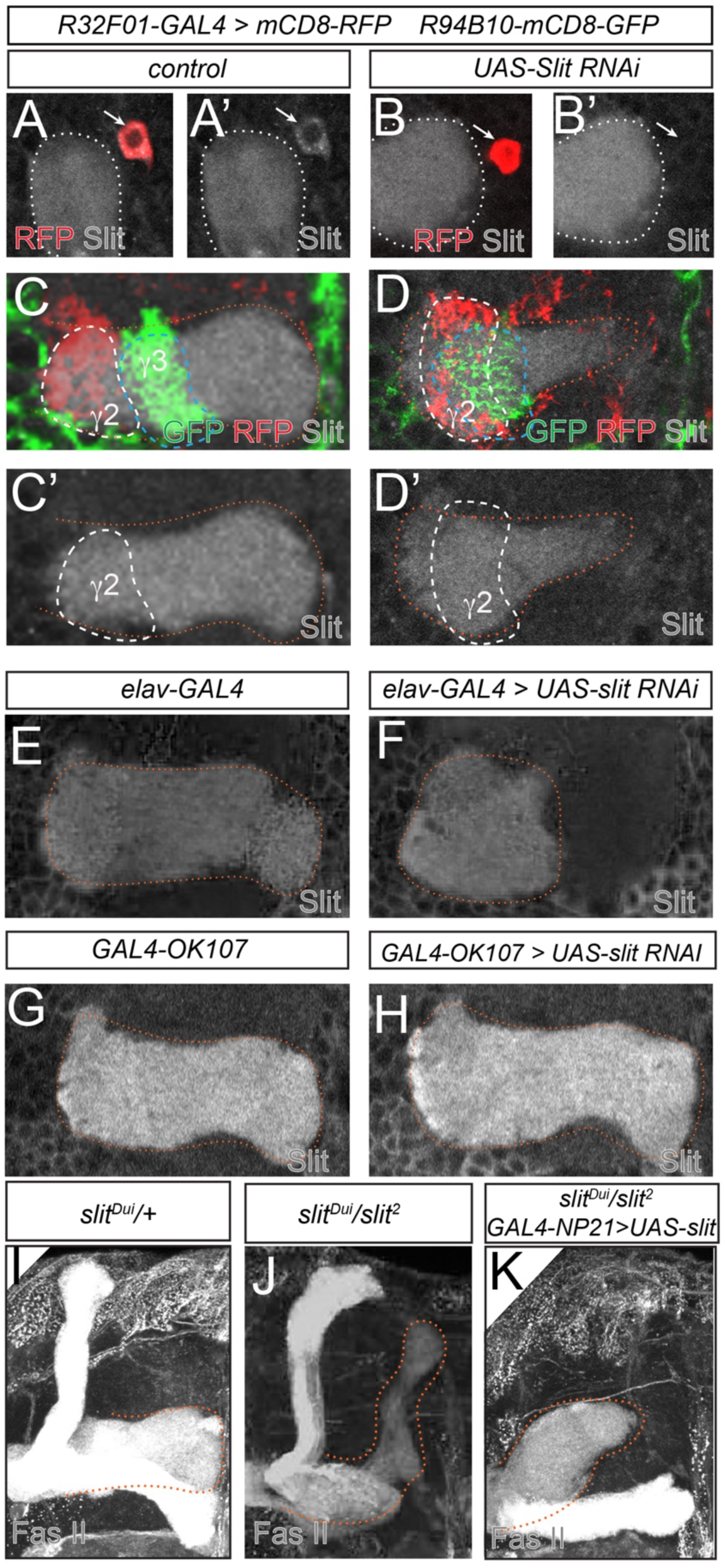
Non-specific binding of anti-Slit antibodies in the MB. (A-B’) Immunostaining with anti-Slit antibodies shows staining signal in the MB calyx (white dotted circles) and the soma of PPL1-γ2 neurons (arrows) (A-A’) and the entire MB γ lobe (B-B’) in a wild type brain. PPL1-γ2 axons (RFP, white dashed circles) and MBON-γ3 dendrites (GFP, blue dashed circles) mark the γ2 and γ3 compartments, respectively. (C-D’) Expressing *UAS-slit RNAi* in PPL1-γ2 neurons diminishes the Slit staining signal in the soma (arrows) of PPL1-γ2 neurons (C-C’) ad leads to projection of MBON-γ3 dendrites (GFP, blue dashed circles) in the γ2 compartment, overlapping with PPL1-γ2 axons (RFP, whit dashed circles), but does not reduce the Slit staining intensity in the γ2 compartment (D-D’). (E-F) Pan-neuronal expression of *UAS-slit RNAi* disrupts the projection of the MB γ lobe (orange dotted circles) but does not reduce the Slit staining intensity in the γ lobe [(F), compared to the wild type in (E)]. (G-H) Expression of *UAS-slit RNAi* in MB neurons neither disrupts the projection of the MB γ lobe (orange dotted circles) nor reduces the Slit staining intensity in the γ lobe [(H), compared to the wild type in (G)]. (I-K) MB γ lobes (FasII staining, orange dotted lines) project normally in *slit^Dui^/+* animals (I) but are mistargeted in *slit^Dui^/slit^2^* mutants (J). Expression of *UAS-slit* in γ neurons does not rescue the mistargeting of the γ lobe in *slit^Dui^/slit^2^* mutants (K).

**Supplemental Fig. S4.**
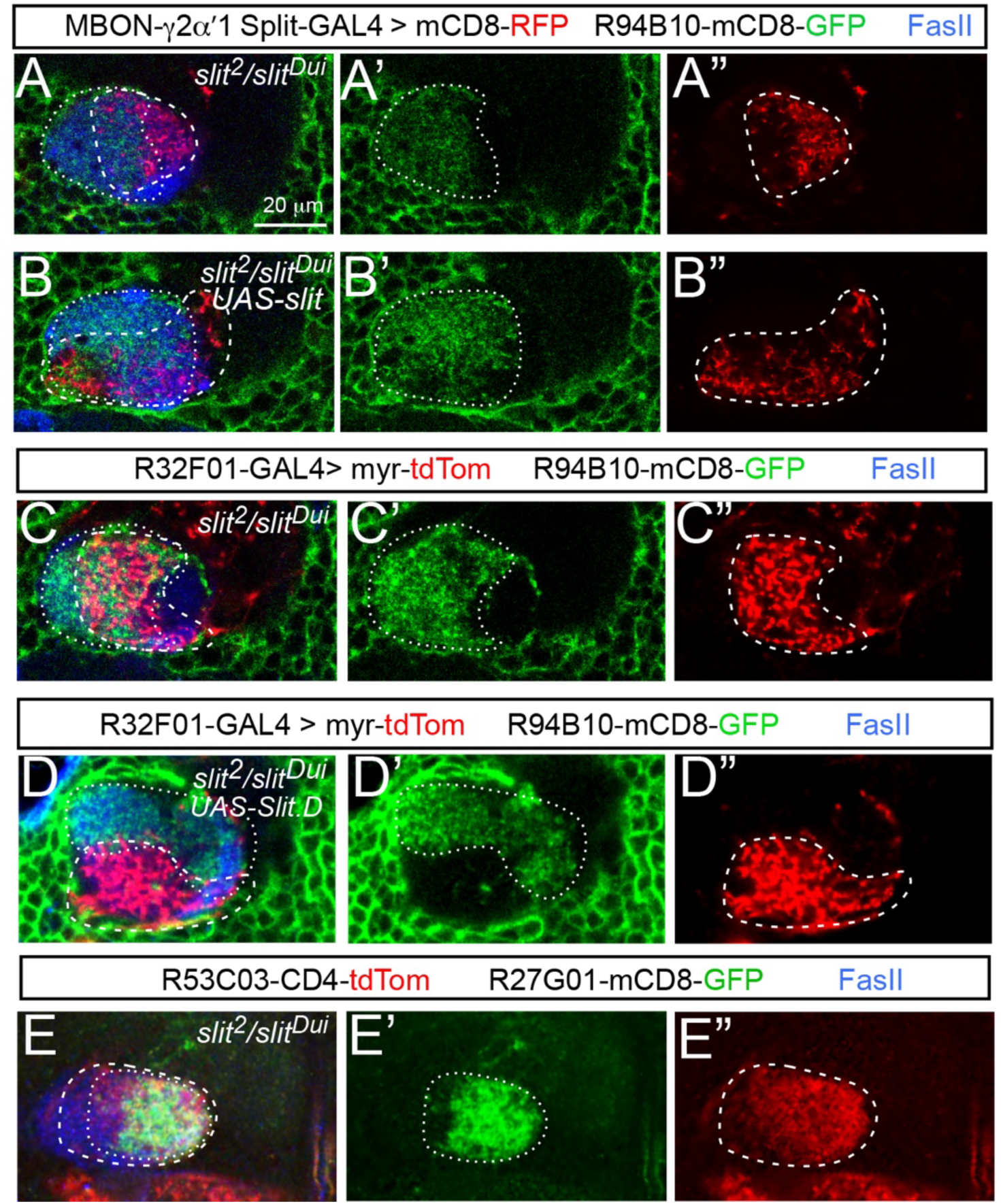
Targeting defects of MBON-γ3 and MBON-γ4 dendrites in *slit* mutants. (A-A”) In a *slit^2^/slit^Dui^* mutant brain, MBON-γ3 dendrites (GFP, dotted circles) project into the γ2 compartment and overlap with MBON-γ2 dendrites (RFP, dashed circles), which are displaced from their original γ2 compartment and shift to the medial side of MBON-γ3 dendrites. (B-B”) Expression of *UAS-Slit* in MBON-γ2 neurons does not restore a sharp boundary between MBON-γ3 dendrites (dotted circles) and MBON-γ2 dendrites (dashed circles), which are displaced from their original γ2 compartment, in *slit* mutants. (C-C”), MBON-γ3 dendrites (GFP, dotted circles) overlap with PPL1-γ2 axons (RFP, dashed circles) in a *slit^2^/slit^Dui^* mutant brain. (D-D”) Expression of *UAS-Slit* in PPL1-γ2 neurons restores a sharp boundary between MBON-γ3 dendrites (dotted circles) and PPL1-γ2 axons (dashed circles) in a *slit^2^/slit^Dui^* mutant brain although their positions in the γ lobe remain altered due to distortion of the γ lobe in *slit* mutants. (E-E”) MBON-γ4 dendrites (tdTom, dashed circles) overlap with MBON-γ5 dendrites (GFP, dotted circles) in a *slit^2^/slit^Dui^* mutant brain. In all images, brains are counterstained with Fas II antibodies to labeled MB γ lobe.

**Supplemental Fig S5.**
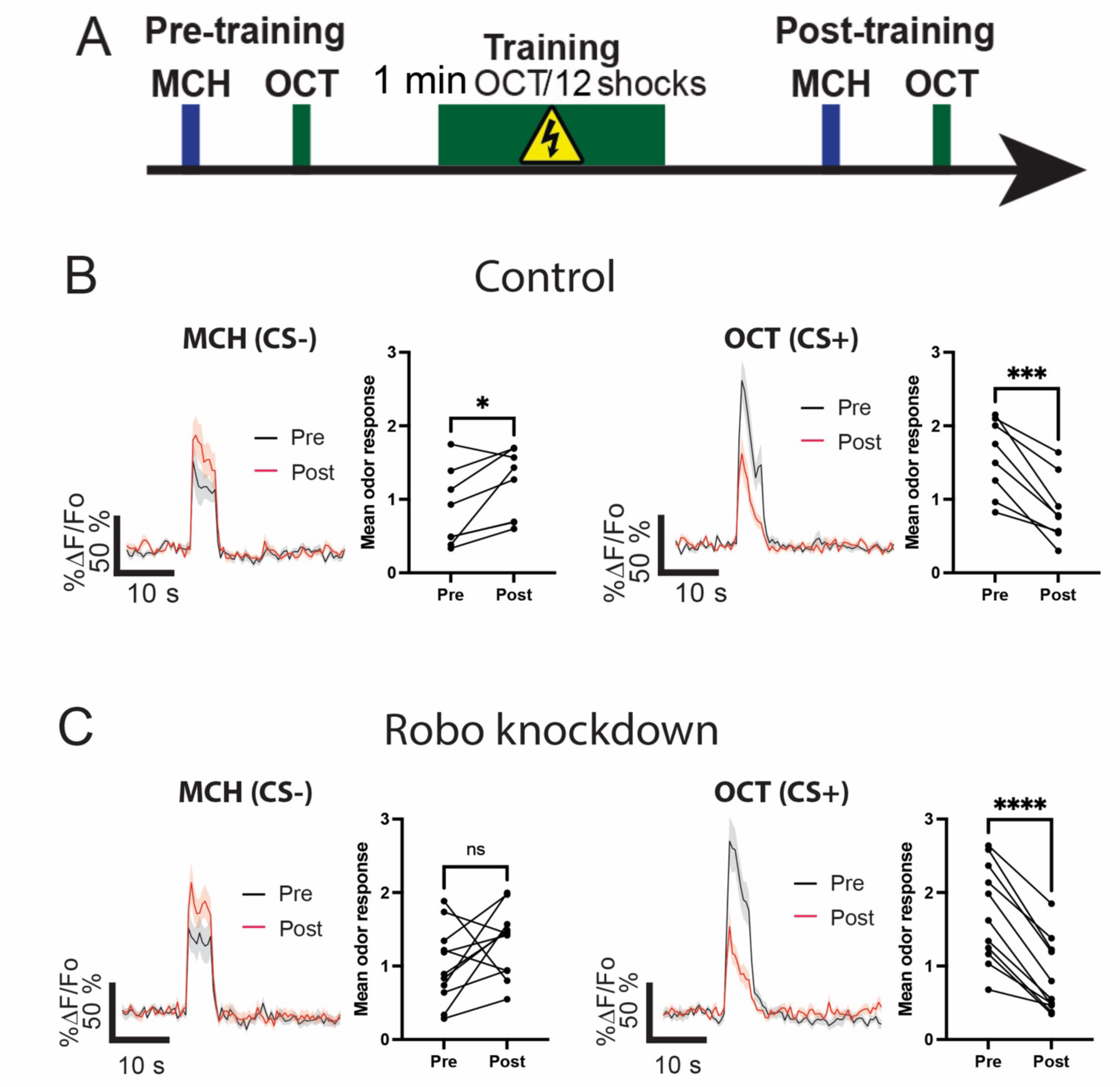
MBON-γ3 neuron memory trace after standard training with 12 electric shocks. (A) Experimental design. Pre-training odor induced calcium response are recorded by presentation of 5-s MCH and OCT with a 30-s ISI; 5 minutes later, flies were aversively trained by pairing 1 min presentation of OCT with 12-90 V shocks. Post-training odor responses were recorded 3 min after. (B-C) GCaMP6f activity in wild type control (A) Robo knockdown (B) MBON-γ3 neuron dendrites in response to MCH (CS−) (left) and OCT (CS+) (right) before and after the standard training with 12 electric shocks.

